# Stathmin-2 loss leads to neurofilament-dependent axonal collapse driving motor and sensory denervation

**DOI:** 10.1101/2022.12.11.519794

**Authors:** Jone Lopez-Erauskin, Mariana Bravo-Hernandez, Maximiliano Presa, Michael W. Baughn, Ze’ev Melamed, Melinda S. Beccari, Ana Rita Agra de Almeida Quadros, Aamir Zuberi, Karen Ling, Oleksandr Platoshyn, Elkin Niño-Jara, I. Sandra Ndayambaje, Olatz Arnold-Garcia, Melissa McAlonis-Downes, Larissa Cabrera, Jonathan W. Artates, Jennifer Ryan, Frank Bennett, Paymaan Jafar-nejad, Frank Rigo, Martin Marsala, Cathleen M. Lutz, Don W. Cleveland, Clotilde Lagier-Tourenne

## Abstract

The human mRNA most affected by TDP-43 loss-of-function is transcribed from the *STMN2* gene and encodes stathmin-2 (also known as SCG10), whose loss is a neurodegenerative disease hallmark. Here using multiple *in vivo* approaches, including transient antisense oligonucleotide (ASO)-mediated suppression, chronic shRNA-mediated depletion in aging mice, and germline deletion, we establish stathmin-2 to be essential for acquisition and maintenance of neurofilament-dependent structuring of axoplasm critical for maintaining diameter and conduction velocity of large-myelinated axons. Sustained stathmin-2 loss from an otherwise mature adult nervous system is demonstrated over a time course of eight months to initiate and drive motor neuron disease that includes 1) shrinkage in inter-neurofilament spacing that is required to produce a three-dimensional space filling array that defines axonal caliber, 2) collapse of mature axonal caliber with tearing of outer myelin layers, 3) reduced conduction velocity, 4) progressive motor and sensory deficits (including reduction of the pain transducing neuropeptide CGRP), and 5) muscle denervation. Demonstration that chronic stathmin-2 reduction is itself sufficient to trigger motor neuron disease reinforces restoration of stathmin-2 as an attractive therapeutic approach for TDP-43-dependent neurodegeneration, including the fatal adult motor neuron disease ALS.

## Introduction

Amyotrophic lateral sclerosis (ALS) is characterized by selective degeneration of upper and lower motor neurons that results in muscle denervation, paralysis, and eventually death typically from respiratory failure within 2-5 years after diagnosis^1^. The loss of neuromuscular junctions (NMJs) is well established as one of the earliest pathological events in both familial and sporadic forms of ALS^2–4^. NMJs are vital synapses formed between motor neuron terminals and muscle cells, and their disruption occurs prior to disease onset and motor neuron degeneration in ALS^3–5^. Importantly, efforts focusing on the maintenance of motor neuron survival have failed to prevent muscle denervation or delay disease onset and progression in mice expressing ALS-causing mutations^6–8^, suggesting that pathological mechanisms involved in muscle denervation occur independently from the death of the motor neuron.

While initial axonal elongation of lower motor neurons can proceed independent of neurofilaments^9^, the initial formation of synapses at NMJs triggers enhanced synthesis and accumulation of neurofilaments^10, 11^, which mediate a >10-fold growth in axonal diameter (and >100-fold increase in axonal volume)^10, 12^. Neurofilaments become the most abundant structural element within mature, myelinated axons, outnumbering the microtubules by a factor of 30 (ref.^13–15^). Importantly, neurofilament pathologies including intracellular inclusions containing neurofilaments have been reported in many neurodegenerative disorders, including ALS^16–19^. Transgenic mice expressing mutated^20^ or altered ratios of neurofilament subunits^21^ develop age-dependent motor neuron disease resembling aspects of human ALS^20, 22^, further emphasizing the importance of neurofilaments in maintaining axonal function. Neurofilament sequence variants have been proposed as contributory risk factors^23, 24^, although mutations in genes encoding neurofilament subunits have not been found as causative of ALS^25^.

The discovery of cytoplasmic mislocalization of the RNA binding protein TDP-43 in affected neurons in >95% instances of ALS and at least 50% of frontotemporal dementia (FTD) profoundly changed the direction of research in ALS and FTD^26, 27^. TDP-43 is known to affect the levels or splicing patterns of mRNAs from 1500 genes^28^. Aside from obvious TDP-43 aggregation, nuclear clearance of TDP-43 has been widely observed in affected neurons in sporadic ALS and FTD^27^ and in the most frequent genetic cause of ALS and FTD, a GGGGCC repeat expansion in the *C9orf72* gene^29–31^. Overwhelming evidence supports that TDP-43 loss of function (and by extension, errors of pre-RNA maturation) are a key aspect of disease mechanism underlying ALS and FTD.

Recently, we and others identified that the stathmin-2 mRNA (encoded by the *STMN2* gene) is the transcript most affected by reduction in TDP-43 function^32, 33^. TDP-43 suppression in human cells drives use of cryptic splice and polyadenylation sites in the *STMN2* pre-mRNA, producing a truncated, non-functional mRNA referred to as exon 2a, that can be found accumulated in brain and spinal cord samples of ALS and FTD patients that develop TDP-43 pathology^32–34^. Importantly, the corresponding loss of stathmin-2 protein (also known as SCG10) results in an impaired ability of human motor neurons to recover after axonal injury^32, 33^. Stathmin-2 mRNAs are among the top 25 most abundant mRNAs in human^32, 35^ and mouse^36^ motor neurons. Notably, TDP-43 dependent regulation of *STMN2* pre-mRNAs found in humans is not conserved in rodents, as the three GU-rich TDP-43 binding motifs and the alternative polyadenylation site in intron 1 of the human *STMN2* gene are not present in the corresponding murine *Stmn2* pre-mRNA^32^. Correspondingly, TDP-43 nuclear clearance or impaired function does not drive loss of stathmin-2 in mice.

Stathmin-2 has been proposed as an important axonal-maintenance factor^37^ and an essential protein for motor axon regeneration^32^. Indeed, after axotomy of sensory axons, stathmin-2 synthesis is upregulated, with the protein accumulating in growth cones^38^. Chronic elimination of stathmin-2 starting from earliest development has been reported to yield mild sensory and motor deficits in young adult mice^39, 40^. Despite an extensive literature on axonal development^41^ and impaired axonal regeneration in injured sensory neurons^37, 38, 42, 43^, it remains unknown whether (and if so how) stathmin-2 contributes to maintenance of motor and sensory axons during aging. Here, we determine that over a time course of months sustained stathmin-2 loss from an otherwise mature adult nervous system provokes shrinkage of neurofilament spacing that defines axonal caliber, axonal collapse with tearing of outer myelin layers, progressive motor and sensory deficits, reduced conduction velocity, and muscle denervation.

## Results

### Denervation and slowed conduction velocity from transient stathmin-2 reduction

Thorough evaluation of accumulated stathmin-2 RNA (determined by *in situ* hybridization — Figure S1a) and protein (determined by immunofluorescence – Figure S1b) within the normal adult nervous system revealed stathmin-2 presence in the soma of virtually every spinal neuron (including interneurons and γ-motor neurons). Stathmin-2 protein also accumulated at the presynaptic side of mature NMJs of gastrocnemius muscle from adult wild type (WT) mice (Figure S1c) and in the outer laminae of the dorsal horn that contains sensory neuron terminals projecting from the adjacent dorsal root ganglion (DRG) (Figure S1b), evidence strongly suggesting a role(s) for stathmin-2 within mature presynaptic motor and sensory neuronal synapses.

Intracerebroventricular (ICV) delivery of a *Stmn2* mRNA-targeting antisense oligonucleotide (ASO) was used to test the consequence(s) of transient suppression of new stathmin-2 synthesis in an otherwise normal adult murine nervous system (Figure 1a). As early as 2 weeks after ASO injection (using either of two different *Stmn2*-targeting ASO variants) into 3-month-old adult mice, *Stmn2* mRNA and protein were suppressed throughout the murine central nervous system (CNS), including cortex (Figure S1d,e) and spinal cord (Figure 1b,S1f). Suppression of *Stmn2* RNAs and accumulated protein were sustained from 2 to 8 weeks post injection in spinal cord (Figure 1c,S1g) and cortex (Figure S1h,i). Although no significant clinical manifestations developed within 8 weeks of lowering accumulated stathmin-2, nerve conduction velocity (the speed at which electrical signals travel along axons) was significantly reduced within the sciatic nerve (Figure 1d) without effect on compound muscle action potential (CMAP) (Figure S1j). Importantly, reduction in stathmin-2 for 8 weeks led to compromised integrity of neuromuscular synapses (Figure 1e), including full denervation of 18% of NMJs and partial denervation of another 29% (Figure 1f).

**Figure 1:**
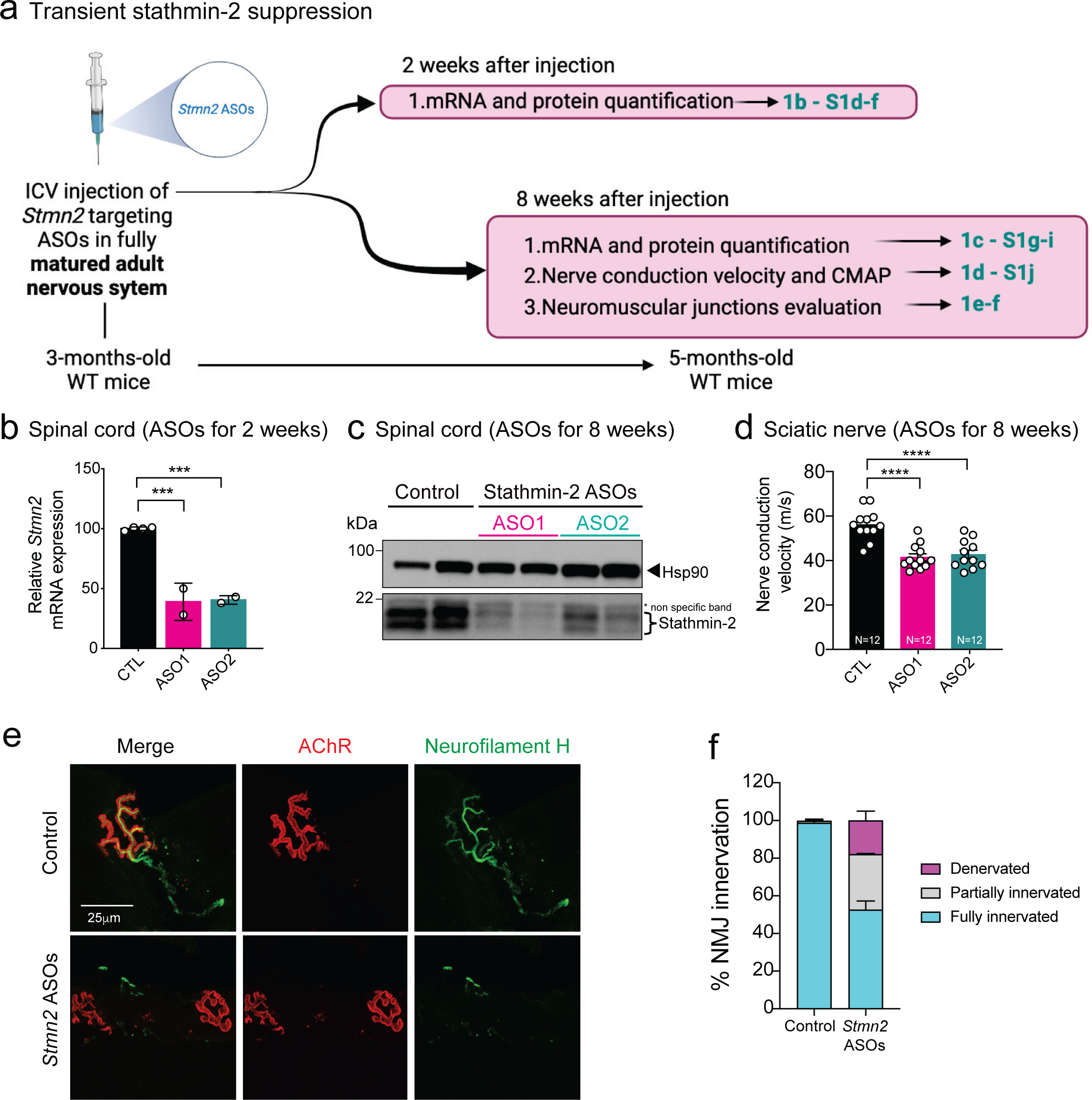
ASO-mediated transient stathmin-2 suppression reduces nerve conduction velocity and triggers muscle denervation. **(a)** Schematic representation of intraventricular (ICV) administration of control or stathmin-2 targeting ASOs (ASO1 and ASO2) in 3 months-old wild-type mice. **(b)** Quantification of *Stmn2* mRNA levels by qRT-PCR 2 weeks after ICV injection. **(c)** Immunoblot showing stathmin-2 protein levels in mice spinal cord extracts 8 weeks after the ICV injection. Hsp90 was used as a loading control. **(d)** Nerve conduction velocity measurement in mice hindlimbs 8 weeks after ICV injections of ASOs. **(e-f)** Representative confocal images **(e)** and innervation rate quantification **(f)** of NMJs in the gastrocnemius muscle 8 weeks after delivery of non-targeting or *Stmn2* targeting ASOs.

### Motor deficits without motor neuron loss from chronic *Stmn2* suppression

The consequence(s) of sustained stathmin-2 loss (thereby mimicking its reduction in ALS) was determined using adeno-associated virus (AAV) to chronically express 1) a microRNA-embedded shRNA^44, 45^ to suppress murine *Stmn2* RNAs in motor and sensory neurons of an otherwise normal adult nervous system (Figure 2a) and 2) a reporter gene encoding mClover (a brighter monomeric GFP derivative) to mark transduced neurons, their axons, and their terminals innervating hindlimb muscles (Figure S2a). Viral vectors encoding shRNA against murine *Stmn2* or an irrelevant, control gene (bacterial β-galactosidase) were introduced into the lumbar spinal cord of 1-year-old WT mice via a single subpial delivery (i.e., injection just beneath the inner-most pia layer^46, 47^). Unlike intrathecal administration, subpial injection in adult animals enables highly effective and long-lasting transduction to deep gray matter neurons and glia of the spinal cord^46, 47^. Additionally, local axonal or synaptic uptake followed by retrograde AAV delivery also transduces cortical motor neurons and sensory neurons of the DRGs. Importantly, lumbar subpial administration achieves efficient focal transduction without targeting cervical or thoracic spinal motor neurons that control forelimbs and respiratory muscles^45^.

**Figure 2:**
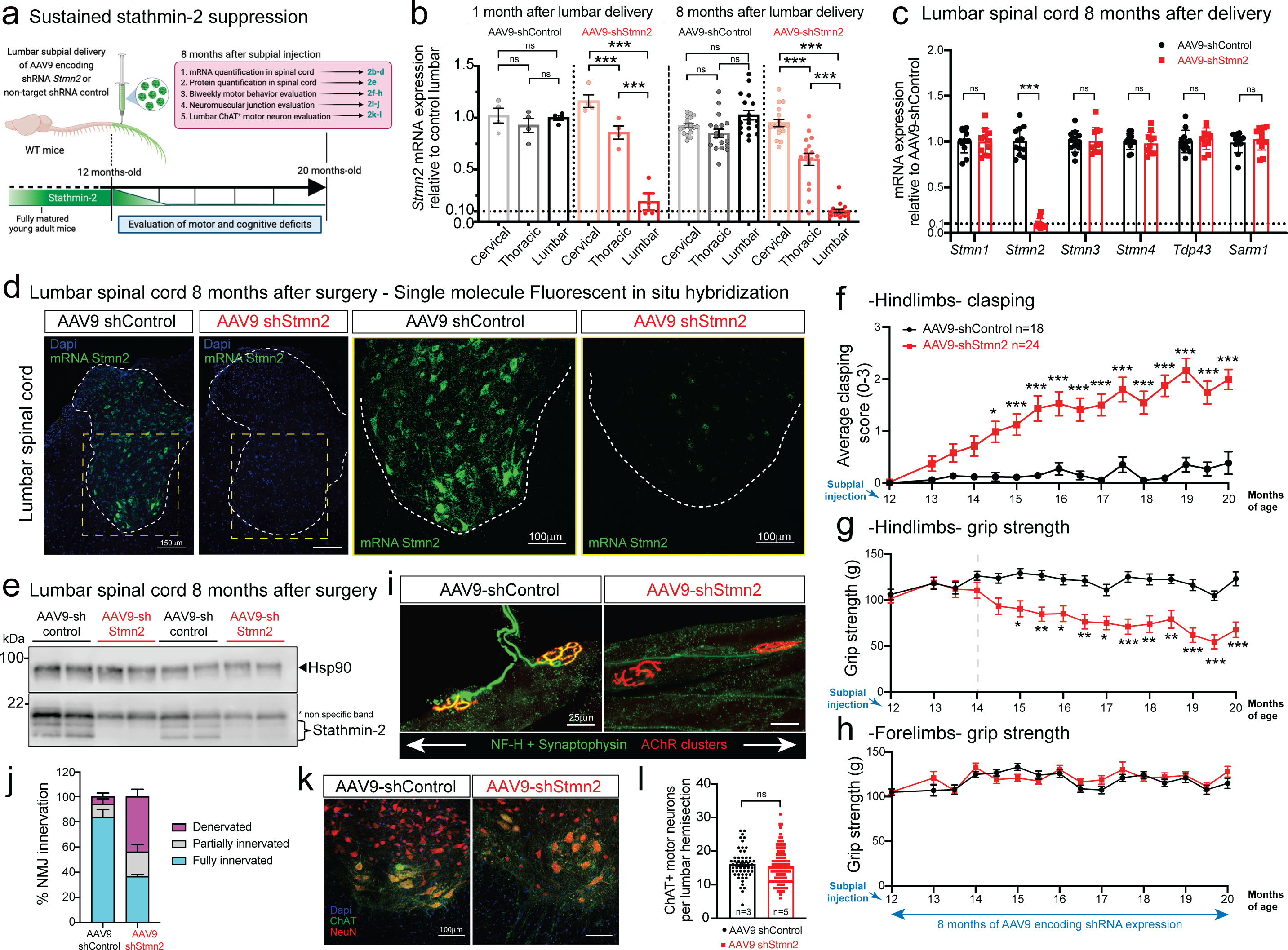
Focal, chronic, and selective suppression of stathmin-2 by subpial delivery of AAV9 shStmn2 in lumbar spinal cord results in motor deficits. **(a)** Schematic illustration of the experimental design for the evaluation of sustained stathmin-2 depletion using AAV9-encoded shRNA subpially injected in 12-month-old adult WT mice. **(b)** Quantification of murine *Stmn2* levels in animals injected with AAV9-encoded shRNA against *Stmn2* or shRNA control across cervical, thoracic, and lumbar segments at 1- and 8-months post-administration. Measurements made on extracted RNA by qRT-PCR using *Gapdh* as an endogenous control gene. Each data point represents an individual mouse. Error bars are plotted as SEM. Statistical analysis by unpaired t-tests ***p<0.001. **(c)** Measurement of mRNA expression by qRT-PCR in lumbar spinal cord segments 8 months after injection of stathmin-2 targeting AAV9 or control. A panel of genes including mouse stathmin1-4 confirms focal and selective *Stmn2* suppression. *Gapdh* was used as an endogenous control gene. Each data point represents an individual mouse. Error bars are plotted as SEM. **(d)** *Stmn2* mRNA levels detected by single molecule FISH in mice lumbar spinal cord. **(e)** Immunoblots of stathmin-2 protein in mice lumbar spinal cord 8 months after subpial injection of AAV9-encoded control or stathmin-2 shRNA. HSP90 was used as a loading control. *Indicates non-specific band. **(f-h)** Longitudinal analysis of hindlimb clasping **(f)**, grip strength **(g)**, and forelimb grip strength **(h)** 8 months after subpial injection of AAV9 encoding a control shRNA (n=18) or shRNA against murine *Stmn2* (n=24); *p<0.05, **p<0.01, ***p<0.001, two-way ANOVA followed by Sidak’s multiple comparison test. **(i)** Representative confocal images of gastrocnemius muscle sections immunostained using synaptophysin and neurofilament-H antibody to mark axon terminals (green) and *α*-bungarotoxin to reveal muscle endplates (red) in 20-month-old mice, 8 months after subpial delivery of AAV9 encoding either non-targeting sequence or shRNA against murine stathmin-2. Scale bar 25*μ*m. **(j)** Quantification of the neuromuscular innervation status. **(k,l)** Representative images **(k)** and quantification **(l)** of ChAT positive motor neurons in the lumbar spinal cord of mice 8 months after subpial delivery of AAV9 encoding either non-targeting sequence or shStmn2.

Analysis of RNAs extracted after subpial administration of AAV9-shStmn2 revealed rapid and sustained reduction (80% at 1 month; 90% at 8 months) of *Stmn2* mRNA selectively within lumbar spinal cord, with only mild or no reduction, respectively, in the thoracic and cervical segments (Figure 2b). There were no compensatory changes in RNAs encoding the other three stathmin genes or other potentially ALS-related genes including TDP-43 and SARM1 or genes whose mRNAs have similar nucleotide sequences that could potentially have been targeted by the shRNA (Figure 2c, S2b). Sustained suppression of *Stmn2* RNAs and resulting protein for 8 months was confirmed with single molecule fluorescence *in situ* hybridization (FISH) (Figure 2d), immunoblotting (Figure 2e,S2c), and immunostaining (Figure S2d). Although body weight over an 8-month course remained unaffected (Figure S2e), within 2 months of reduced stathmin-2 synthesis, motor performance was impaired, including induction of hindlimb clasping (Figure 2f) and reduced hindlimb grip strength (Figure 2g), accompanied by loss of muscle volume as observed in the gastrocnemius muscle (Figure S2f). Accordingly, biweekly analyses revealed that progressive lumbar motor deficiencies developed up to 20 months of age (Figure 2f,g) without effect(s) on cervical spinal cord motor function (Figure 2h).

Sustained stathmin-2 suppression resulted in disruption of more than half of the NMJs of lumbar motor neurons, with 43% of total (70% of affected) NMJs fully denervated after 8 months of stathmin-2 reduction (Figure 2i,j). Despite this, there was no reduction in the number of lumbar ChAT+ motor neurons (Figures 2k,l) and no sign of neuroinflammation measured by changes in number or morphologies of astrocytes or microglia in the lumbar spinal cord (Figure S2g-i). Thus, sustained loss of stathmin-2 in motor neurons of an otherwise fully mature, adult nervous system drives muscle denervation without motor neuron loss.

### Sustained loss of stathmin-2 provokes neurofilament-mediated axonal collapse

Continued suppression of stathmin-2 synthesis within lumbar spinal motor neurons of otherwise normal adult WT mice yielded a 35% reduction in conduction velocity of the sciatic nerve (Figure 3a), similar to the conduction velocity reduction reported in ALS patients^48–50^. Conduction velocity is thought to be determined by two primary factors: the degree of myelination and axonal diameter, with bigger diameters driving faster conduction velocity^51, 52^. Recognizing this, measurement of axonal diameters following 8-month suppression of stathmin-2 synthesis revealed a significant shrinkage in cross sectional area of the axons of the α-motor neurons (with diameters > 3μm) responsible for muscle contraction and a corresponding shrinkage in γ-motor axons (diameters < 3μm). The average α-motor axonal diameter was reduced from 7.5μm to 5.5μm (Figure 3b,c), corresponding to a ∼50% reduction in cross sectional area and producing the expected overall shrinkage (by 29%) of lumbar motor axon roots (Figure 3d), while the number of γ- and α-motor axons remained unchanged (Figure 3e).

**Figure 3:**
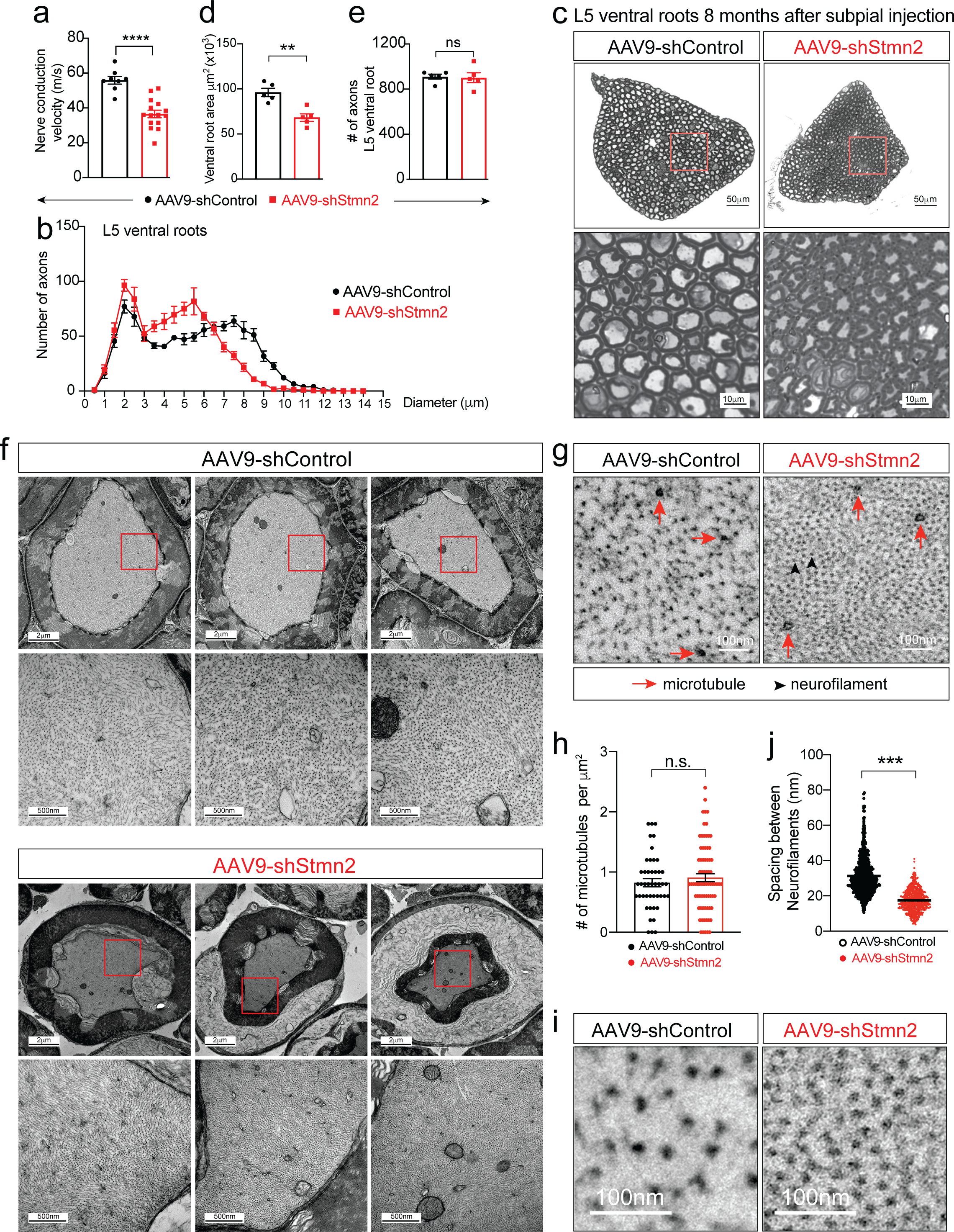
Sustained loss of stathmin-2 reduces nerve conduction velocity and provokes axonal collapse by decreasing the spacing between axonal neurofilaments. **(a)** Nerve conduction velocity on 20-month-old WT mice 8 months after lumbar subpial delivery of control or *Stmn2* shRNAs. Statistical analysis by unpaired t-tests ****p<0.0001. **(b)** Size distribution of motor axon diameters in the L5 motor roots from 20-month-old WT mice 8 months after subpial injection of AAV9 encoding non-targeting shRNA or *Stmn2* targeting shRNA. Error bars are plotted as SEM. **(c)** Representative micrographs of motor roots and higher magnification images of ventral root motor axon morphology and diameters, 8 months after subpial injection of AAV9 encoding non-targeting shRNA or *Stmn2* targeting shRNA. **(d-e)** Quantification of cross-sectional area **(d)** and of the total number of axons per ventral root **(e)**. Each data point represents an individual mouse. Error bars are plotted as SEM. Statistical analysis by unpaired t-tests **p<0.01. **(f)** Representative electron microscopy images of large caliber axons in the motor roots and increased magnification micrographs of the axolemma showing altered spacing between neurofilament filaments in WT mice 8 months after subpial injection of AAV9-sh*Stmn2* when compared to AAV9-shControl. Scale bar 2*μ*m for upper panels, and 500nm for lower panels. **(g)** Representative images of ventral root axoplasm where neurofilaments (black arrowheads) outnumber microtubules (red arrows). **(h)** Number intra-axonal microtubules per *μ*m^2^ of ventral root axoplasm. **(i-j)** Representative images **(i)** and quantification of spacing distance between neurofilaments in the axoplasm of each group **(j)**. Statistical analysis by unpaired t-tests ***p<0.001.

Sustained reduction in stathmin-2 also produced dramatic changes in axoplasm and the interaction of the axon with its myelinating Schwann cells (Figure 3f). Electron microscopic imaging of α-motor axons revealed obvious axonal shrinkage, with the innermost myelin layers remaining attached to the axon, but with shrinkage-induced tearing within the outermost myelin layers, thereby at least partially disrupting connection of the Schwan cell body to its encapsulated axon. Axoplasm of the shrunken axons appeared collapsed with compartments of highly compacted neurofilaments (Figure 3f). Although overall microtubule number was unchanged (Figure 3g,h), neurofilament organization was highly abnormal, with average interfilament spacing reduced by nearly half (from 31.2 nm to 17.4 nm – Figure 3i,j).

Neurofilaments are responsible for the acquisition of normal axonal caliber^13, 53–56^ and are assembled as obligate heteropolymers^57^ of three highly conserved polypeptide subunits: neurofilament light (NF-L), medium (NF-M), and heavy (NF-H). In mice, the long tails of NF-M and NF-H contain 7 or 51 lysine-serine-proline (KSP) phosphorylation sites, respectively, that extend from the filament surface to determine mean interfilament spacing^56^ and interactions with other cytoskeletal components and organelles^56^. A 10-20-fold increase in axonal caliber (corresponding to a 100-400 increase in axonal volume) initiates concomitantly with myelination that occurs within the first three postnatal weeks. An initial burst in caliber occurs in the first two to three months and is dependent on an “outside-in” signaling cascade from the myelinating cell to the underlying axon that requires the tail domain of NF-M^56^.

While levels of the three neurofilament subunits remained unchanged following sustained stathmin-2 suppression within the spinal cord (Figure S3a-e), levels of phosphorylated NF-H and NF-M were reduced (Figure S3f-h), the latter of which has been shown in mice to be required for establishing normal axonal caliber^56^. These results reveal a crucial role of stathmin-2 in the maintenance of axoplasm through orchestrating the “outside-in” phosphorylations of the larger two neurofilament subunits that act to mediate their nearest neighbor distances and the assembly of a three-dimensional, space-filling, interlinked neurofilament array that determines axonal caliber.

### Reduced stathmin-2 levels impair tactile perception

Retrograde delivery of an AAV9 encoding GFP was used to validate that subpial injection yielded transduction of the majority of sensory neuron soma residing in the adjacent DRGs (Figure 4a,S4a), consistent with prior reports^45, 58^. Virtually all normal DRG neurons express *Stmn2* mRNAs detectable by FISH, albeit at diverse levels (Figure 4b,c). Eight months after subpial administration of AAV9-shStmn2, *Stmn2* mRNAs (detected by single molecule FISH – Figure 4b) and stathmin-2 protein (detected by indirect immunofluorescence – Figure 4d) were reduced in the majority of DRG neurons, with more than half expressing murine *Stmn2* mRNA at ∼30% of initial level (Figure 4b,c) and a corresponding reduction in detectable stathmin-2 protein (Figure 4d). These results were consistent with an overall 53% reduction of the initial *Stmn2* mRNA level (measured by qRT-PCR in mRNA extracted from lumbar DRGs – Figure 4e).

**Figure 4:**
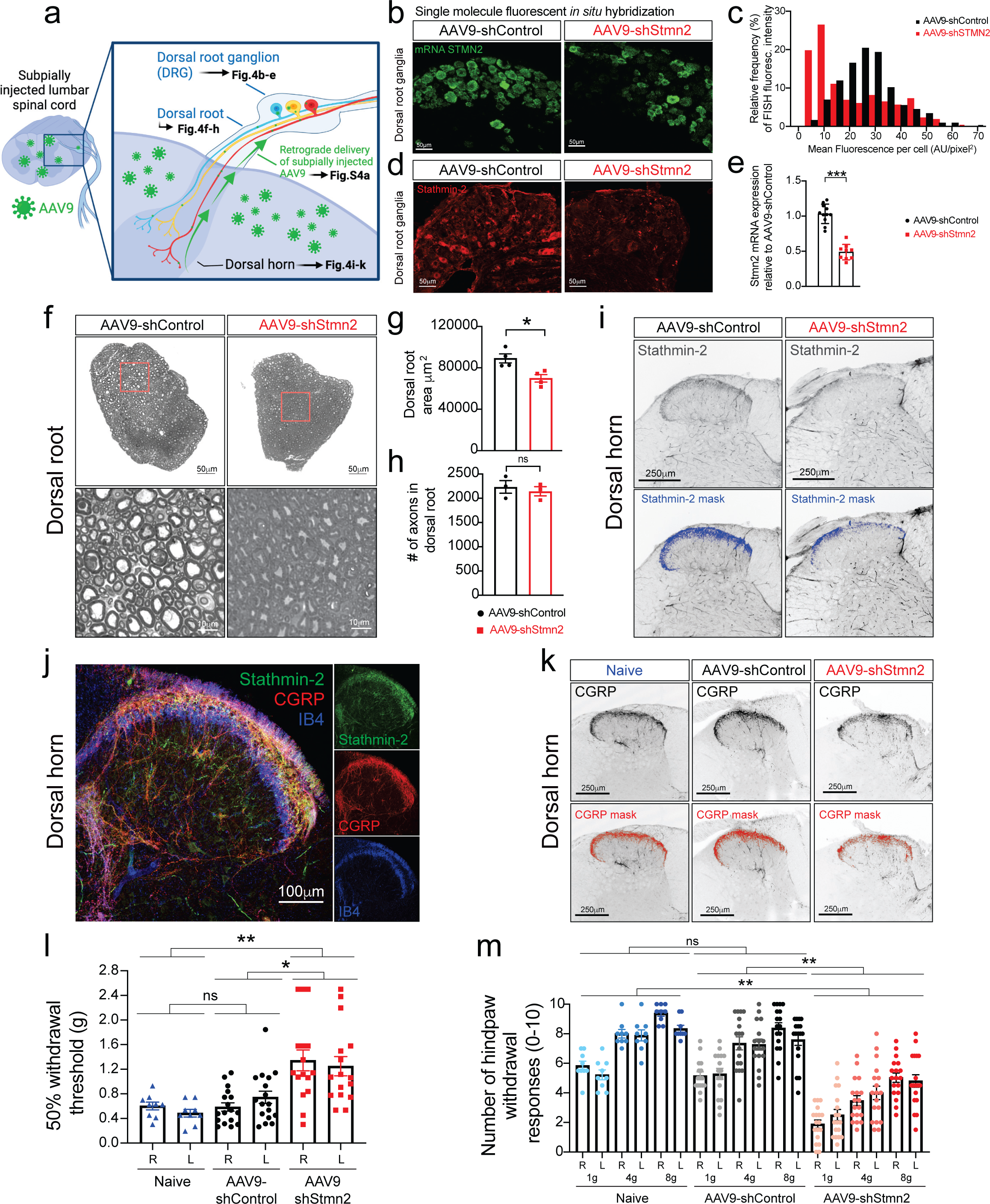
Reduced stathmin-2 levels in lumbar dorsal root ganglia impair hindlimb sensory system. **(a)** Schematic of the strategy to determine the impact of subpially injected AAV9-encoded shRNA on the pseudo-bipolar sensory neurons located in the dorsal root ganglion innervating the dorsal spinal cord. **(b-d)** Representative confocal images showing *Stmn2* mRNA levels by single molecule FISH **(b)** and its fluorescence distribution **(c)**, and stathmin-2 protein **(d)** in the lumbar dorsal root ganglion 8 months after the subpial administration of AAV9 encoding either non-targeting or *Stmn2* targeting shRNA. **(e)** Quantification of *Stmn2* mRNA levels in dorsal root ganglion 8 months after the subpial administration of AAV9 encoding either non-targeting or *Stmn2* targeting shRNA. *Gapdh* is used as an endogenous control gene. Each data point represents an individual mouse. Error bars plotted as SEM. Statistical analysis by unpaired t-tests ***p<0.001. **(f-h)** Representative images of entire dorsal roots and higher magnification images showing axonal morphology and diameter size **(f)**, quantification of dorsal roots crossectional area **(g)**, and total number of sensory axons **(h)** on cross-sectioned dorsal root of WT mice, 8 months after subpial administration of AAV9 encoding either control or stathmin-2 targeting shRNA. **(i)** Representative dorsal horns of lumbar spinal cord sections immunolabeled with stathmin-2 antibody at 20 months of age, 8 months after subpial administration of either non-targeting shRNA or *Stmn2* targeting shRNA. Blue mask represents stathmin-2 signal in dorsal horn. **(j)** Representative confocal micrograph with WT dorsal horn immunostained with stathmin-2, CGRP, and IB4. **(k)** Representative dorsal horn areas of lumbar spinal cord sections immunolabeled with CGRP antibody. Red mask represents CGRP positive signal in dorsal horn of lumbar spinal cords in 20 month old, age matched non-injected (naive) mice, or 8-months after subpial delivery of either non-targeting shRNA or *Stmn2* targeting shRNA. **(l)** Quantification of the 50% withdrawal threshold upon von Frey filament based on mechanical stimuli of mice hindlimbs at 20 months of age in non-injected mice or mice 8 months after subpial injection with AAV9 encoding non-targeting shRNA or *Stmn2* targeting shRNA. Each data point represents an individual animal. Error bars are plotted as SEM. Statistical analysis by one-way ANOVA Tukey’s multiple comparison test, *p<0.05 **p<0.01. **(m)** Quantification of hind paw response to increasing von Frey filament force stimuli. Each data point represents an individual animal. Error bars are plotted as SEM. Statistical analysis by one-way ANOVA Tukey’s multiple comparison test, *p<0.05 **p<0.01.

Sensory neurons extend an axon that bifurcates, with one branch directed towards the periphery to sense environmental stimuli, while the other branch enters the dorsal root to innervate neurons of the dorsal spinal cord (Figure 4a)^59^. As observed in ventral roots (Figure 3b,c), the size distribution of axons in the dorsal root shifted towards smaller diameters upon sustained reduction in synthesis of stathmin-2 (Figure S4b,c), producing an overall 22% shrinkage of cross-sectional area of the affected dorsal root (Figure 4f,g) without any axonal loss (Figure 4h). Reduction of stathmin-2 was also observed in the sensory neuron terminals innervating the dorsal horn of the lumbar spinal cord of AAV9-shSTMN2 mice (Figure 4i, S4d), with the area occupied by stathmin-2 positive terminals diminished by two-thirds within 8-months of sustained suppression (Figure S4d). Interestingly, the stathmin-2-containing sensory terminal area in the dorsal horn colocalized with calcitonin gene-related peptide (CGRP)-positive fibers (Figure 4j) corresponding to unmyelinated, slowly conducting peptidergic C fibers (Lamina I) and thinly myelinated more rapidly conducting Aγ fibers (outer Lamina II) (reviewed in ref^59–,61^). By contrast, the inner lamina II mainly comprised of unmyelinated, slowly conducting non-peptidergic C fibers (identifiable by isolectin B4 (IB4) staining) barely overlapped with stathmin-2 positive terminals (Figure 4j), indicating specificity of stathmin-2 role in peptidergic terminals of pain transmitting sensory neurons.

Consistently, loss of stathmin-2 in the sensory terminals was accompanied by a reduction of CGRP positive fibers innervating the dorsal spinal cord (Figures 4k, S4e), evidence of the importance of stathmin-2 in terminal maintenance of peptidergic sensory neurons. These molecular alterations in sensory neurons altered somatosensory behavior. Mice with sustained reduction of stathmin-2 had suppressed tactile evoked responses (i.e., increased paw withdrawal thresholds) when compared to mice injected with non-targeting AAV9-shControl (Figure 4l) or age-matching non-injected, naïve animals. Response to tactile stimuli was also impaired in response to mechanical nociceptive stimulus (pain). Indeed, while naïve and AAV9-shControl animals behaved indistinguishably, responses to nociceptive mechanical stimuli (determined by hindpaw withdrawal frequency upon increasing force) were reduced in mice with diminished stathmin-2 (Figure 4m).

### Loss of stathmin-2 in embryogenesis leads to increased perinatal lethality

CRISPR/Cas9 genome editing was used to inactivate one endogenous *Stmn2* allele in C57BL/6J mice by deletion of a 1028bp segment including exon 3 (Figure S5a). The deletion was predicted to produce an unstable RNA that is a substrate for nonsense-mediated decay as a consequence of a frameshift mutation at codon 38 (Figure S5a). Although mice were born in expected Mendelian ratios from breeding *Stmn2*^+/-^ animals (Figure S5b), 80% of *Stmn2*^-/-^ pups died shortly after birth (median survival of only 1.5 days), indicating one or more roles for stathmin-2 in early development. However, in *Stmn2*^-/-^ mice that survived to weaning age, no further accelerated lethality was observed (Figure 5a). As expected, stathmin-2-encoding mRNAs were reduced or undetectable, respectively, in cortical (Figure 5b) or spinal cord (Figure 5c) RNAs from *Stmn2*^+/-^ or *Stmn2*^-/-^ adult mice. Stathmin-2 protein was also reduced by 50% in heterozygous *Stmn2*^+/-^ mice and was undetectable in *Stmn2*^-/-^ mice in lumbar spinal cord protein extracts (Figure 5d, S5c) and with immunostaining of ventral motor neurons in lumbar spinal cord sections (Figure 5e). Reduction or complete absence of stathmin-2 did not produce compensatory changes in expression of the other stathmin family members (stathmin-1, stathmin-3, and stathmin-4) (Figure S5d, S5e).

**Figure 5:**
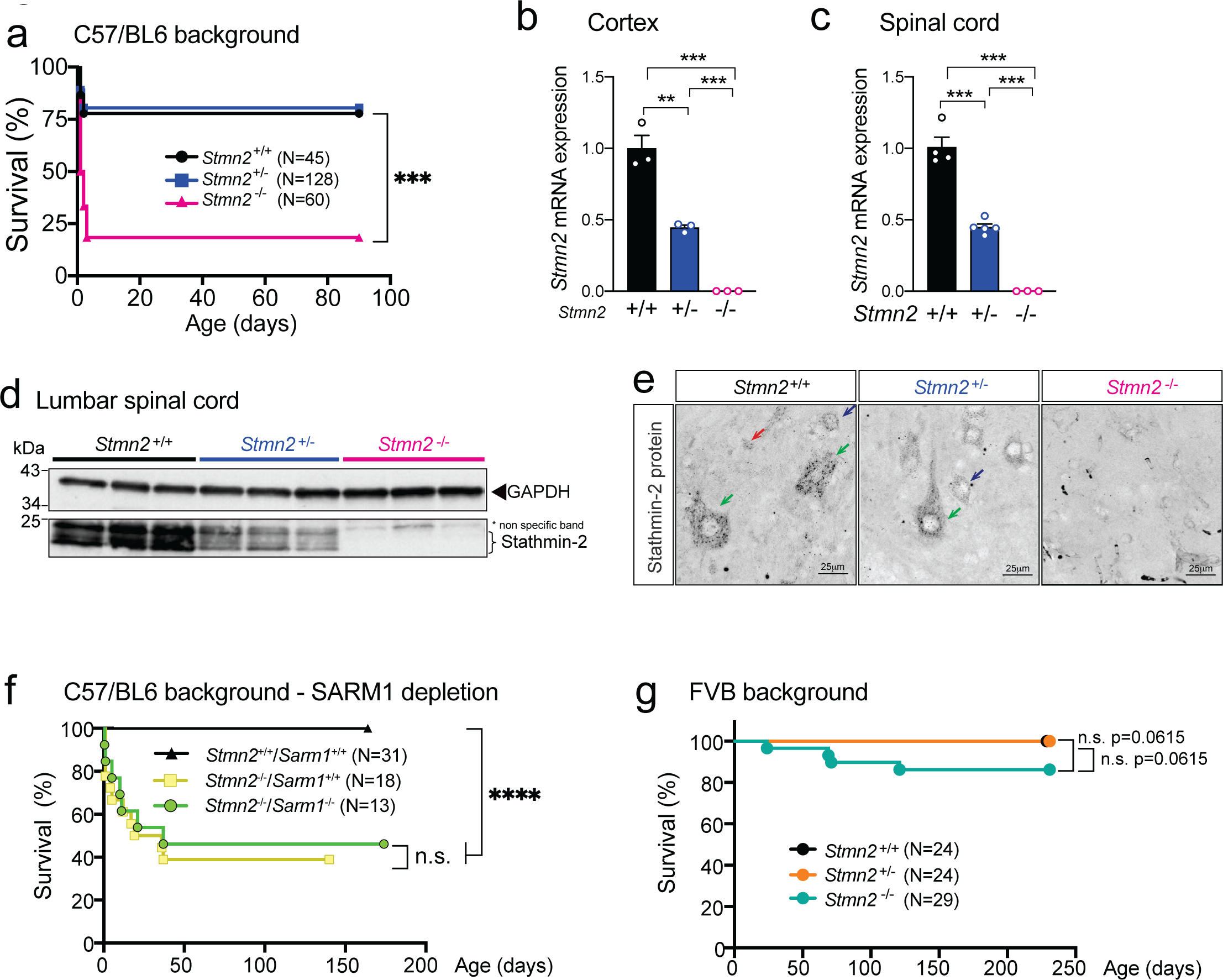
Stathmin-2 has an important role early after birth rescued by FVB genetic background but not by *Sarm1* ablation. **(a)** Survival curve of *Stmn2*^+/+^, *Stmn2*^+/-^ and *Stmn2*^-/-^ mice in a C57/BL6 background. **(b-c)** Measurement of murine *Stmn2* mRNA levels extracted from cortex **(b)** and spinal cord **(c)** of *Stmn2*^+/+^, *Stmn2*^+/-^ and *Stmn2*^-/-^ mice. *Gapdh* was used as an endogenous control gene. Each data point represents an individual mouse. Error bars are plotted as SEM. Statistical analysis by unpaired t-tests **p<0.01 ***p<0.001 **(d)** Immunoblot showing levels of the ∼21 kD mouse stathmin-2 protein from 3 different animals per genotype. GAPDH was used as a loading control. **(e)** Confocal micrographs of stathmin-2 immunolabeling at the ventral spinal cord of 12-month-old *Stmn2*^+/+^, *Stmn2*^+/-^ and *Stmn2*^-/-^ mice. Green arrows: *a*-motor neurons; blue arrows: *y*-motor neurons; red arrows: interneurons. **(f)** Survival curve of *Stmn2*^+/+^/*Sarm1*^+/+^, *Stmn2*^-/-^/*Sarm1*^+/+^ and *Stmn2*^-/-^ /*Sarm1*^-/-^ mice in a C57/BL6 background. Statistical analysis by Log-rank Mantel-Cox Test ****p<0.0001. **(g)** Survival curve of *Stmn2*^+/+^, *Stmn2*^+/-^ and *Stmn2*^-/-^ mice in FVB background. Statistical analysis by Log-rank Mantel-Cox Test.

The threshold of stathmin-2 levels needed to rescue perinatal lethality in *Stmn2*^-/-^ mice was determined by breeding with two newly generated BAC transgenic lines harboring a complete human *STMN2* gene. Expression of human stathmin-2 at 10% of the endogenous protein level (Figure S5f – corresponding to BAC mouse line 9446 with 1 copy of the transgene – Figure S5g) was not enough to counteract the lethality after birth (8% of progeny that survived to p10 were *Stmn2^-/-^*, BAC^9446^, only 1/3^rd^ of the 25% expected). However, lethality was fully rescued (24% of the progeny survived at p10 from 25% expected) by accumulation of BAC-encoded human stathmin-2 to 50% of the endogenous protein level (Figure S5h – corresponding to BAC mouse line 9439 with 3 copies of the transgene – Figure S5g).

### Stathmin-2 function during development is independent of the SARM1 pathway

Both stathmin-2 and NMNAT2 (nicotinamide mononucleotide adenylyltransferase 2 which functions in NAD+ biosynthesis) have been proposed as axonal survival factors^62, 63^ that act to oppose the toll-like receptor adapter <u>s</u>terile <u>a</u>lpha and <u>T</u>IR <u>m</u>otif containing <u>1</u> (SARM1). SARM1 is implicated as a central executioner of axonal degeneration (referred to as Wallerian degeneration) and its suppression has been proposed as a therapeutic strategy for multiple neurodegenerative diseases, including ALS^62, 63^. While loss of *Nmnat2* induces perinatal lethality that is restored by ablation of *Sarm1*^64^, perinatal death driven by stathmin-2 loss was unaffected by reduction in or absence of *Sarm1* in cohorts of *Stmn2*^-/-^ mice bred to have neither or both *Sarm1* alleles inactivated (Figure 5f). Thus, postnatal death from *Stmn2* loss is independent of SARM1 pro-degeneration activity.

### Perinatal lethality from stathmin-2 loss is mouse strain and environment dependent

Inbred genetic backgrounds can significantly influence expression of phenotypes associated with known genetic perturbations^65^ and can underlie variation in disease severity between individuals with the same mutation(s). To test if this was true for stathmin-2, we backcrossed the B6-*Stmn2*^+/-^ mice into an alternative genetic background, the FVB mouse strain. In contrast with the widespread perinatal lethality of *Stmn2*^-/-^ mice in the C57BL/6J background, FVB:B6 mice deficient for *Stmn2* were born in the expected Mendelian ratios without significant perinatal lethality (Figure 5g). Added to this, the absence of perinatal lethality in an additional *Stmn2*^-/-^ mouse line in the C57BL/6J background^39^ contrasts with the perinatal lethality seen in our C57BL/6J *Stmn2*^-/-^ mice (in colonies maintained at Jax or UCSD) and in the colony of DiAntonio and colleagues^40^, indicating that there must be as yet unidentified genetic and environmental factors that influence stathmin-2’s contribution to perinatal survival.

### Absence of stathmin-2 results in motor deficits and muscle denervation without motor neuron loss

The effects of homozygous *Stmn2* loss from earliest development were determined in FVB mice and in C57BL/6J mice that survived to weaning age. Body weight in males (Figure 6a,b) and females (Figure 6c,d) was reduced compared to age-matched *Stmn2*^+/-^ and WT littermates. Similar to the motor impairment developed following chronic AAV9-shStmn2 suppression in the normal adult nervous system, motor performance [evaluated by hindlimb clasping (Figure 6e,f), grip strength (Figure 6g,h), and rotarod (Figure 6i,j)] was significantly impaired at 3 months of age in both *Stmn2*^-/-^ mice strains compared to WT mice. Once developed, initial motor deficits in *Stmn2^-/-^* mice did not progress after 3 months of age. Motor performance of *Stmn2*^+/-^ mice was indistinguishable from WT littermates.

**Figure 6:**
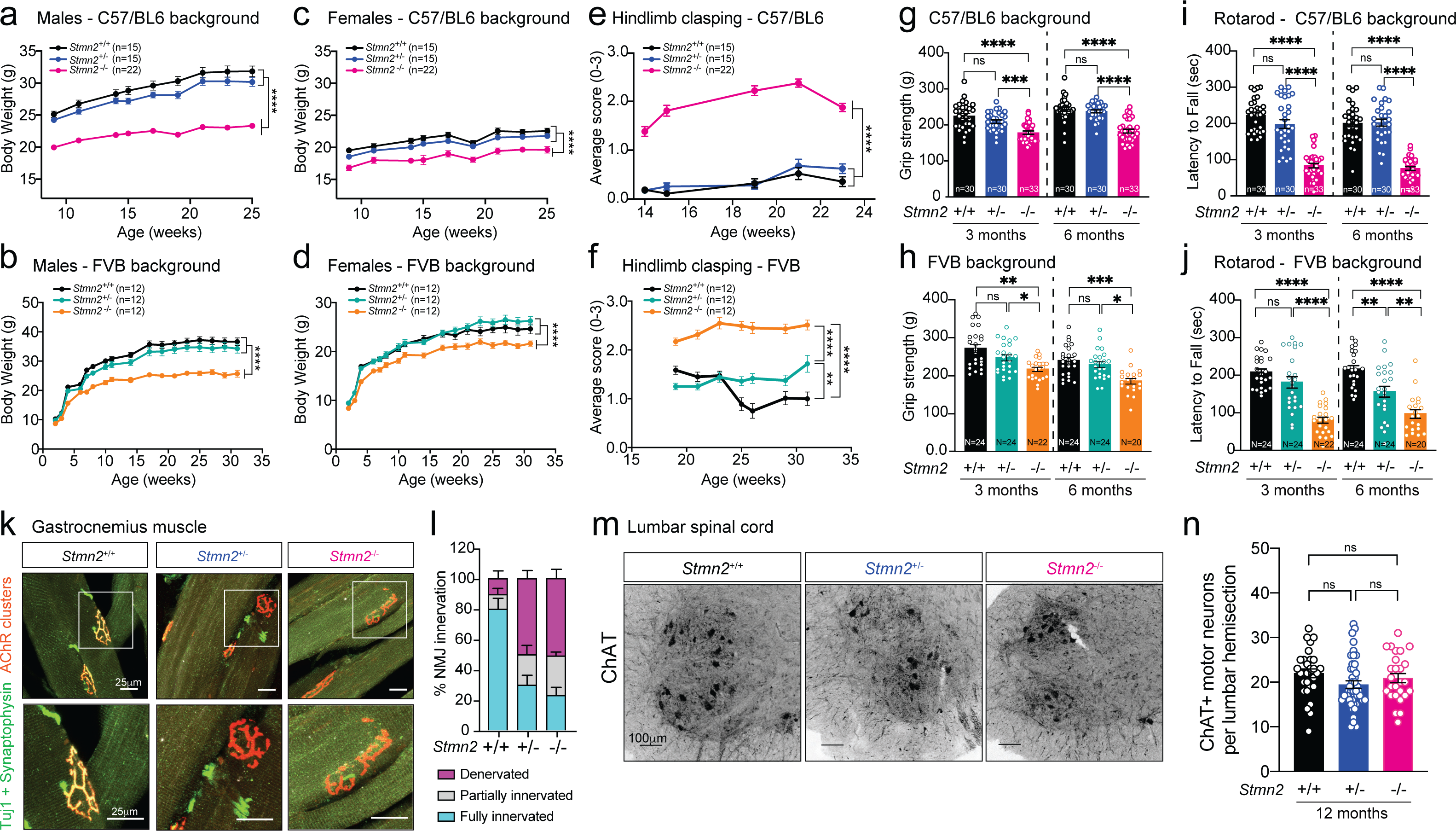
Absence of stathmin-2 results in motor deficits and muscle denervation without motor neuron loss. **(a-b)** Body weight from *Stmn2*^+/+^, *Stmn2*^+/-^ and *Stmn2*^-/-^ male mice in the C57/BL6 background **(a)** and FVB background **(b)**. **(c-d)** Body weight from *Stmn2*^+/+^, *Stmn2*^+/-^ and *Stmn2*^-/-^ female mice in the C57/BL6 **(c)** and FVB **(d)** backgrounds. **(e-f)** Hindlimb clasping measurements of *Stmn2*^+/+^, *Stmn2*^+/-^ and *Stmn2*^-/-^ mice in the C57/BL6 **(e)** and FVB **(f)** backgrounds. **(g-h)** Hindlimb grip strength measurements of *Stmn2*^+/+^, *Stmn2*^+/-^ and *Stmn2*^-/-^ mice at 3 and 6 months of age in the C57/BL6 **(g)** or FVB **(h)** backgrounds. **(i-j)** Rotarod performance of *Stmn2*^+/+^, *Stmn2*^+/-^ and *Stmn2*^-/-^ mice at 3 and 6 months of age in the C57/BL6 **(i)** or FVB **(j)** backgrounds. **(k)** Gastrocnemius muscle sections from *Stmn2*^+/+^, *Stmn2*^+/-^ and *Stmn2*^-/-^ mice with axon terminals immunolabelled using a combination of synaptophysin and beta-III-tubulin (Tuj1) antibodies (green) and muscle endplates using *α*-bungarotoxin (red). **(l)** Innervation frequency quantified in gastrocnemius muscles immunolabeled with axon terminal and muscle endplates markers and quantified using confocal microscopy imaging. **(m)** Representative lumbar spinal cord sections of 12-month-old *Stmn2*^+/+^, *Stmn2*^+/-^ and *Stmn2*^-/-^ mice motor neurons immunolabelled with ChAT antibody. **(n)** ChAT positive neurons quantified in the ventral spinal cord of lumbar hemisections.

Loss of stathmin-2 from early embryogenesis in *Stmn2^-/-^* mice induced NMJ denervation of adult hindlimb muscles comparable to that following 8 months of AAV9-shStmn2-mediated reduction of stathmin-2 within a fully matured motor neuron (Figure 6k,l), accompanied by mildly reduced CMAP when compared to aged-matched WT and *Stmn2*^+/-^ mice (Figure S6a). Denervation was not accompanied by a reduction in the number of ChAT positive spinal motor neurons even at 12 months of age (Figure 6m,n). Once again, presence or absence of a functional SARM1-dependent degenerative pathway did not affect development of motor deficits upon loss of stathmin-2 (Figure S6b). Thus, even when stathmin-2 loss is initiated in earliest embryogenesis, the resultant alterations in distal axons that trigger conduction velocity and motor deficits are independent of SARM1 activity and are insufficient to compromise adult motor neuron survival. In addition, examination of *Stmn2*^-/-^ mice confirmed reduced nociception relative to *Stmn2*^+/+^ mice (Figure S6c), accompanied by reduced accumulation of CGRP (Figure S6d), findings reinforcing the importance of stathmin-2 in adult sensory systems.

### Absence of stathmin-2 inhibits radial axonal growth and nerve conduction velocity

Complete absence of stathmin-2 yielded reduced conduction velocity (in comparison to WT littermates) in the sciatic nerves of 3- and 6-month-old C57BL/6J *Stmn2*^-/-^ mice (Figure 7a). Measurement of axonal diameters at 3 months of age in L5 ventral motor axons of *Stmn2*^-/-^ mice revealed reduced diameters (Figure 7b,c) without any axonal loss (Figure 7d), demonstrating inhibited or delayed axonal caliber acquisition, a feature that is normally completed in mice by 3 months of age^66^. Levels of all three neurofilament subunits (and their KSP phosphorylated forms) were comparable in *Stmn2*^-/-^ and *Stmn2*^+/+^ mice at 3 months of age (Figure S7a-f). By 12 months of age, however, absence of stathmin-2 yielded markedly diminished levels of NF-H and its phosphorylated form (pNF-H, detected with the SMI-31 monoclonal antibody) (Figure 7e,f, quantified in Figure 7g,h), as well as reduced levels of NF-M (Figure 7i; quantified in Figure 7j) and its phosphorylated form (pNF-M, also detected with SMI-31 antibody in Figure 7e and quantified in Figure 7l). No changes were observed in NF-L levels among the genotypes at the different timepoints tested (Figures 7k,m and S7a,e). Thus, absence of stathmin-2 impaired accumulation of NF-M and NF-H, as well as their phosphorylation, which together leads to an expected reduced acquisition or maintenance of axonal calibers^56, 67–69^ and provides a mechanistic explanation for reduced nerve conduction velocity^70^.

**Figure 7:**
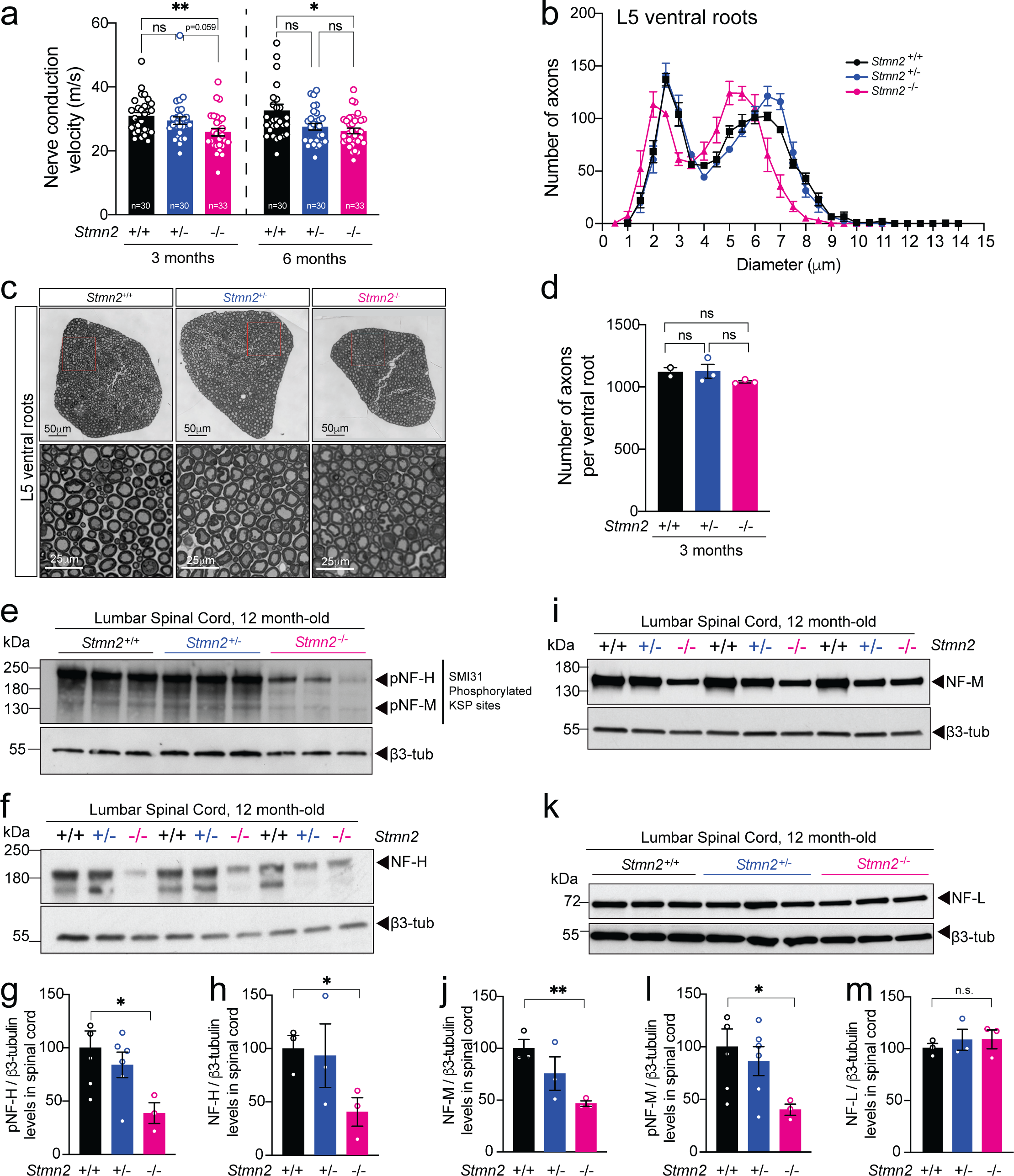
Absence of stathmin-2 reduces nerve conduction velocity and alters axonal radial growth, and neurofilament composition. **(a)** Nerve conduction velocity of *Stmn2*^+/+^, *Stmn2*^+/-^ and *Stmn2*^-/-^ mice at 3 and 6 month of age in the C57/BL6 background. **(b)** Size distribution of motor axons in the L5 ventral motor roots of 3-month-old *Stmn2*^+/+^, *Stmn2*^+/-^ and *Stmn2*^-/-^ mice in the C57/BL6 background. **(c)** Representative motor roots micrographs and higher magnification images from 3-month-old *Stmn2*^+/+^, *Stmn2*^+/-^ and *Stmn2*^-/-^ mice in the C57/BL6 background showing reduced axon diameter in *Stmn2*^-/-^. **(d)** Number of axons quantified in *Stmn2*^+/+^, *Stmn2*^+/-^ and *Stmn2*^-/-^ mice L5 ventral roots. **(e,f)** Immunoblotting for phosphorylated forms of neurofilament heavy (pNF-H) and neurofilament medium (pNF-M) **(e)** and total neurofilament heavy (NF-H) **(f)** in spinal cord protein extracts of 12 months-old *Stmn2*^+/+^, *Stmn2*^+/-^ and *Stmn2*^-/-^ mice. **(g-i)** Quantifications of pNF-H **(g)**, NF-H **(h)**, and pNF-M **(i)** normalized to the amount of β3-tubulin from immunoblotting in **(e)** and **(f)** respectively. β3-tubulin remained unchanged upon the same amount of protein loading, thus it was used as a loading control. **(j-m)** Immunoblotting for neurofilament medium (NF-M) **(j)** and neurofilament light (NF-L) **(k)** in spinal cord protein extracts of 12 months-old *Stmn2*^+/+^, *Stmn2*^+/-^ and *Stmn2*^-/-^ mice. Quantification for NF-M **(l)** and NF-L **(m)** normalized to the amount of β3-tubulin are shown.

## Discussion

Using transient or sustained suppression, we have established the importance of stathmin-2 in acquisition and maintenance of proper caliber of motor axons, in neurofilament-dependent structuring of axoplasm, in acquisition and retention of normal conduction velocity, and in maintenance of mature NMJs and CGRP^+^ sensory terminals. Sustained reduction of stathmin-2 (to 10% of the initial level) in an otherwise mature nervous system induces collapse of mature axonal caliber which underlies the tearing of outer myelin layers, thereby at least partially disconnecting the “outside-in” signaling cascade from myelinating cell to the ensheathed axon^56^. Correspondingly, neurofilament phosphorylation is inhibited, resulting in the collapse of the inter-neurofilament spacing that is required to produce the three-dimensional space filling array that mediates axonal caliber^56^. This in turn yields motor neuron disease that includes reduced conduction velocity, progressive motor and sensory deficits (including diminished tactile and nociceptive somatosensory responses), and NMJ denervation. Even transient reduction of stathmin-2 in the adult nervous system is sufficient to provoke denervation of NMJs.

Chronic reduction of stathmin-2 alone initiates distal motor axonal degeneration independent of the SARM1 prodegenerative pathway, including failure of synaptic maintenance and obvious axonal shrinkage, with axons peeling away from the layers of close packed myelin and axoplasmic collapse into compartments of highly disorganized, closely spaced neurofilaments. Although stathmin-2 has been previously proposed to regulate the rapid growth and shrinkage of microtubules (referred to as dynamic instability)^71–74^ through its direct binding to α/β tubulin heterodimers^41^ in a phosphorylation-dependent manner^75^, our data establish that reduction in stathmin-2 in a normal adult nervous system provokes strikingly altered neurofilament organization without affecting microtubule number. Reduced level and phosphorylation of NF-M and NF-H upon stathmin-2 loss reflect disruption of the “outside-in” signaling pathway from myelinating cell to underlying axoplasm that is required to mediate and maintain axonal diameter^56^.

Perhaps most importantly, chronic reduction of stathmin-2 in motor neurons in an otherwise normal adult nervous system is itself sufficient to drive the initial steps of an ALS-like motor phenotype, a finding that strongly supports a mechanistic contribution to disease initiation and progression from the established stathmin-2 loss in sporadic and familial ALS pathogenesis^35^. Although TDP-43 loss of function affects the levels or splicing of more than 1500 RNAs^28, 76^, the mRNA encoding stathmin-2 is the transcript most affected in human neurons and its restoration alone is sufficient to rescue compromised axonal regeneration after injury of TDP-43 depleted motor neurons^32, 33^. Beyond motor neurons, our effort has also identified a significant role of stathmin-2 in tactile and nociceptive pain transmission and expands restoration or maintenance of stathmin-2 as a therapeutic target for sensory neuropathies [e.g., facial onset sensory and motor neuropathy (FOSMN)^77–79^] that include TDP-43 mislocalization and aggregation in sensory neurons.

In addition to its requirement for axoplasmic and synaptic maintenance in the adult, our evidence establishes that absence of stathmin-2 in mice can dramatically compromise perinatal survival (in the C57/BL6J background) independent of the SARM1 prodegenerative pathway, consistent with an independent study^40^. The near absence of early lethality in a different genetic background (FVB) implicates contribution(s) of yet unidentified genetic variants to perinatal survival in the absence of stathmin-2. Added to this, unidentified environmental factors can also be determinants of stathmin-2 loss-dependent perinatal death, since housing of C57/BL6J *Stmn2*^-/-^ mice in a different vivarium^39^ can mitigate the lethality seen in our colonies (housed at Jax or UCSD), similar to prior reports for C9-ALS/FTD disease mouse models^80^.

Finally, we have demonstrated that not only does stathmin-2 accumulate at the axon terminals of mature motor and sensory neurons in the adult nervous system (e.g., NMJs and dorsal horn lamina I and outer laminal II, respectively), but also that there is a continuing requirement for stathmin-2 in the maintenance of motor and sensory neuron synapses. Upon sustained reduction in stathmin-2, synapses are disrupted, triggering a “dying back” degenerative process similar to that recognized as an early event in ALS^3^. However, reduction in stathmin-2 does not compromise motor neuron survival in mice, at least within the 8-month timeframe we have analyzed. This might be due to additional mRNA alterations occurring in TDP-43 proteinopathies, including the loss of *UNC13A*^81, 82^ that may compound the pathology driven by loss of stathmin-2 alone. Nevertheless, our data strongly support stathmin-2 loss as an early contributor to ALS disease initiation and progression and highlight the attractiveness of restoring stathmin-2 as a therapeutic approach for TDP-43-dependent neurodegenerative diseases.

## Methods

### Animals

Female and male adult C57BL/6 mice (∼12 months of age) were obtained from the Cleveland Laboratory (University of California San Diego (UCSD)). *Sarm1* deficient mice (B6.129X1-*Sarm1^tm1Aidi^*/J, catalog number 18069) were obtained from The Jackson Laboratory. The general health and body weight of all the animals were monitored on a regular daily basis during the whole experiment. All experiments were performed in accordance with the National Institutes of Health (NIH) Guidelines and approved by the Institutional Animal Care and Use Committee (IACUC) and animal care and use program (ACP) at UCSD and IACUC-committee at The Jackson Laboratory.

#### Generation of Stmn2 deficient mice

*Stmn2* deficient mice were generated by targeted-deletion of *Stmn2*-exon 3 using CRISPR/Cas9 technology. Two RNA guides were used to target sites upstream (Stmn2_up_crRNA1: TGCGCAGACTCCATCAGACT; Stmn2_up_crRNA2: ATTTTACACTCTGCTCTATG) and downstream (Stmn2_down_crRNA1: ATCCTACTGTAGAGAATTGA; Stmn2_down_crRNA2: ACTATGGACATTAAGACTGG) of exon 3 in the *Stmn2* gene. We identified by sequencing a founder mouse carrying a 1028 nucleotide deletion spanning *Stmn2*-exon 3. This deletion is designed to cause a change of amino acid sequence after residue 38 and an early truncation 46 amino acids later. This mouse was backcrossed to C57BL/6 by two generations and maintained as heterozygous to establish a stable stock (C57BL/6J-Stmn2^em2Lutzy^/Mmjax, here B6-*Stmn2^(+/-)^*, catalog number JR33740). Expression of 50% of stathmin-2 protein was confirmed in brain and spinal cord of heterozygous *Stmn2*^+/-^ mice and was undetected in *Stmn2*^-/-^ mice (Figure 1). For mixed genetic background experiments, B6-*Stmn2^(-/+)^*, were outcrossed to the strain FVB/NJ (catalog number JR1800) for one generation and heterozygous F1’s were intercrossed to generate F2 FVB;B6-*Stmn2^(-/-)^* homozygous mice.

#### Generation of human BAC STMN2 transgenic lines

We generated a BAC transgenic mouse in C57BL/6J background by microinjecting the BAC clone RRP11-761J24, a 202,106bp DNA fragment carrying the complete human STMN2 gene, into zygotes derived from C57BL/6J mice. A transgenic founder was identified, and stock established (catalog number JR36029). For genetic rescue experiments B6-*Stmn2^(-/+)^* were crossed to B6-*Tg (STMN2)*. F1 heterozygous *Stmn2^(-/+)^* and hemizygous *Tg(STMN2)* mice were crossed back with B6-*Stmn2^(-/+)^* to generate B6-*Stmn2^(-/-)^Tg(STMN2)* (catalog number JR 36033 and 36034), homozygous for the mouse *Stmn2* null allele and hemizygous for the human BAC transgene. The numbers of progeny mice generated that were homozygous for the Stmn2 KO mutation and carriers or non-carriers for each of the four transgenic lines were determined in at least 150 progeny per strain to determine if the presence of the human STMN2 transgene could rescue the 80% perinatal lethality associated with loss of murine Stmn2 on the C57BL/6J genetic background. JR 36033 line containing BAC transgenic lines 9739 demonstrated rescue of the murine lethality phenotype. The remaining strain, JR 36034 demonstrated no BAC-dependent survival. All strains were genotyped by PCR as containing both the human STMN2 exon 1 and exon 5 regions.

### Human STMN2 copy number analysis on BAC-Tg lines

Transgene copy number was analyzed by droplet digital PCR (ddPCR) on genomic DNA (gDNA) samples of hemizygous mice. Briefly, 20 ng of gDNA were used by reaction. A FAM-labeled probe was used to target human STMN2 gene (dHsaCNS772300147, Bio-Rad) and a Hex-labeled probe was used to target mouse ApoB (dMmuCNS407594696, Bio-Rad) as two copies reference gene. Copy number was determined using Quantasoft application from Bio-Rad and reported as copies/mouse genome.

### RNAi AAV vector development

One hundred Psm2 97-mer hairpin oligos were designed against the 3’UTR of the murine STMN2 mRNA using the “Hannon Lab RNAi Central siRNA design tool” (http://katahdin.mssm.edu/siRNA/RNAi.cgi?type=shRNA). Top candidate hairpin sequences were hand-picked using internal sequence content criteria and bioinformatic filtering to eliminate potential off-targets. The top five hairpin design sequences produced as ssDNA oligonucleotides (IDT) for further development and amplified as previously described^44^ for plasmid integration into a miR30a backbone vector by restriction digest and ligation. A control hairpin was similarly produced targeting the bacterial beta-galactosidase gene (with no predicted human or mouse target). The resulting microRNA-embedded RNAi elements were subcloned into a custom AAV transfer vector encoding an RNA Polymerase II driven human Ubiquitin C promoter and Clover fluorescent protein coding sequence, with the RNAi elements inserted between the clover stop codon and an SV40 polyadenylation signal sequence. All bacterial plasmids were grown in One Shot Stbl3 chemically competent E. coli (Thermo) at 30°C to prevent recombination of hairpin sequence or ITR elements. Candidate and control vectors were tested for activity by transient transfection with Trans-IT LT1 transfection reagent (Mirus) into murine neuron-like N2A cells followed by Trizol RNA isolation (Thermo) and qRT-PCR quantification with primers amplifying murine *Stmn2* and *Gapdh* mRNAs. A GFP-only transfection was used as a comparative control. The vector with the top activity against murine *Stmn2* was produced at high volume alongside the control vector for viral packaging (>400ug plasmid per transfer vector), followed by complete sanger sequencing vector verification in 1% DMSO containing reactions. AAV9 rep/cap and helper plasmids were produced in parallel, and virus was generated at the UCSD viral core by 3-plasmid transfection into 293T cells, with virions purified as previously described^46^. Viral genome titers were measured by limiting dilution series and qPCR using primers to amplify the Clover coding sequence and using the transfer plasmid dilution series as a quantification reference. Hairpin oligonucleotide sequences are listed in (Table 1).

**Table 1.**
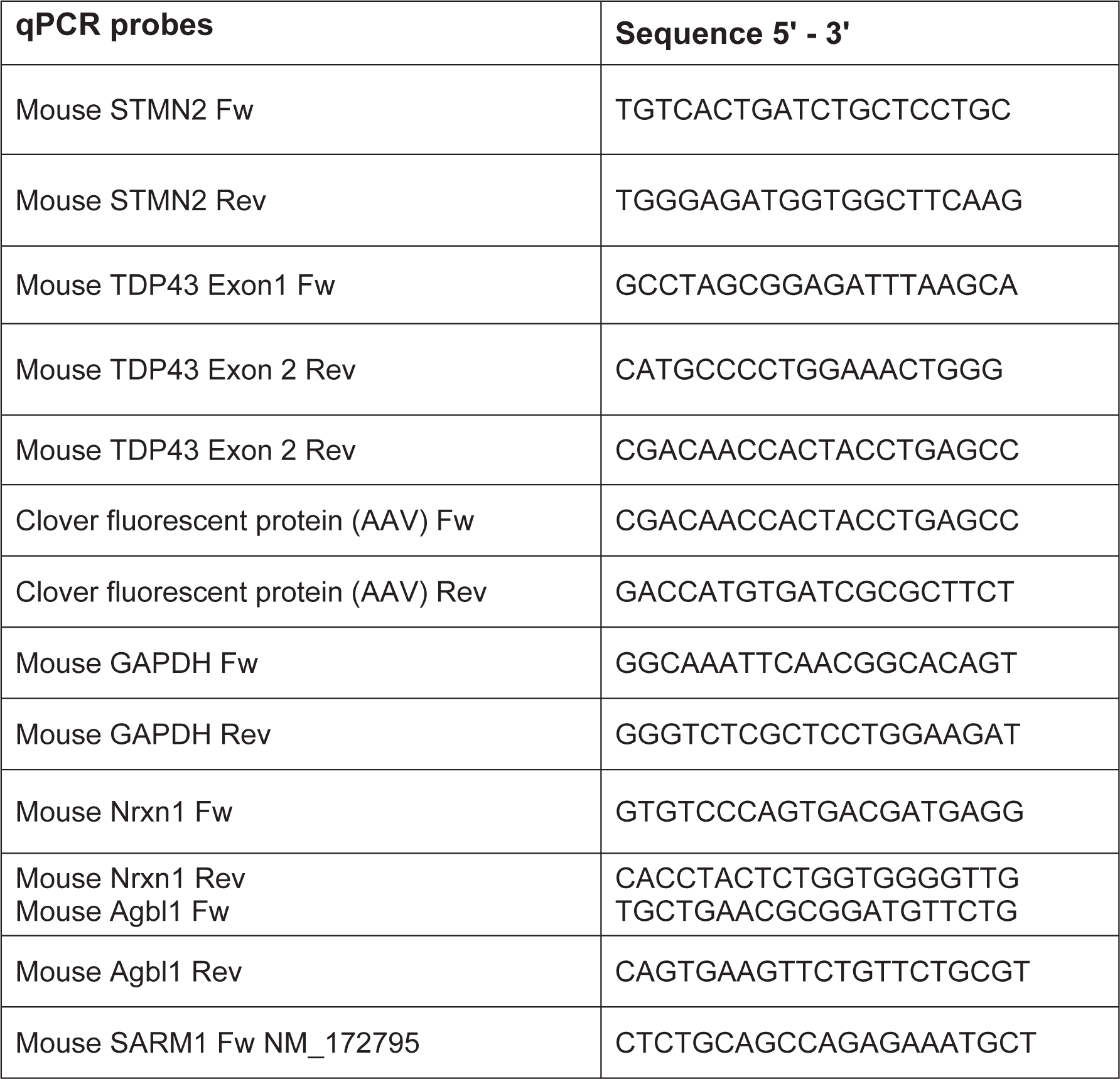
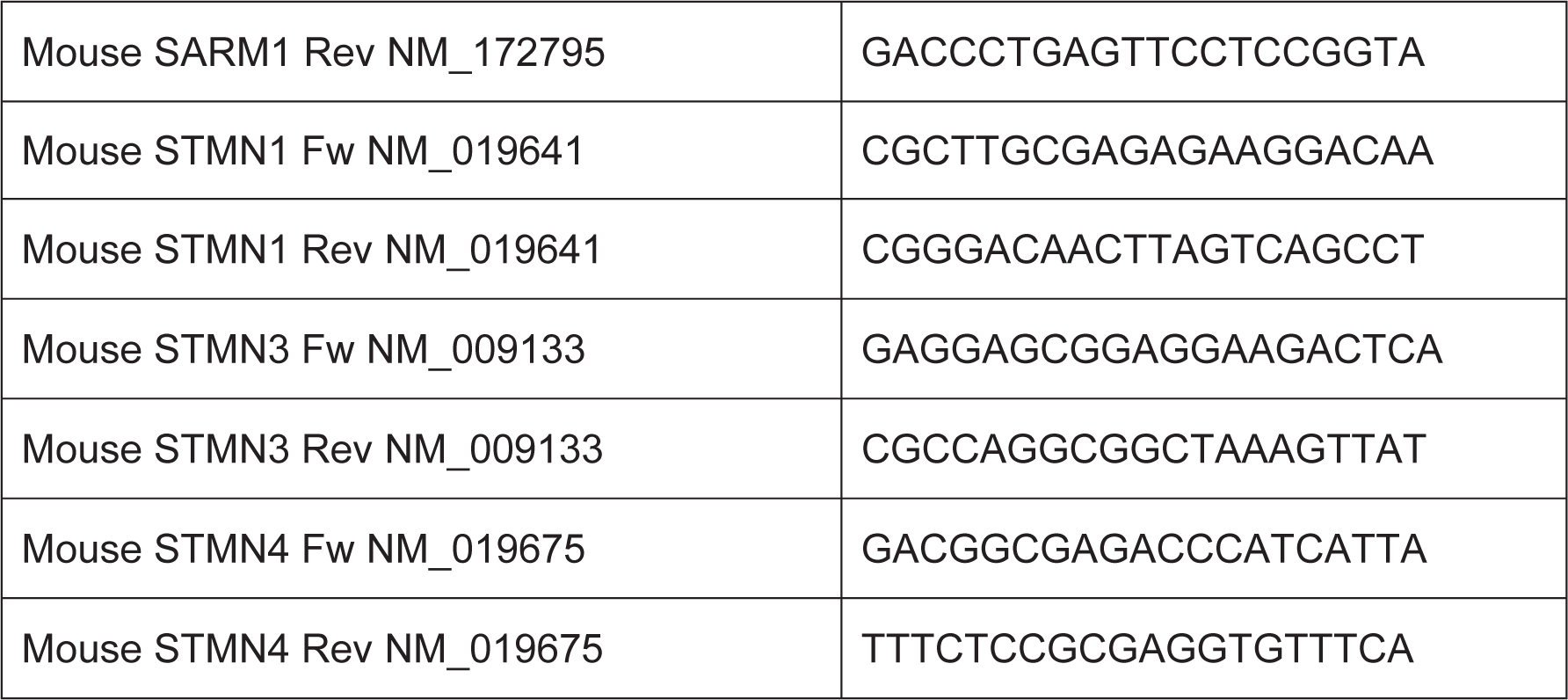
qPCR Probes.

### Subpial injections

Surgical aseptic technique was the standard in every procedure. Subpial vector delivery in mice was performed as previously described^83^. Briefly, in anesthetized mice with Isofluorane 2-3%, a skin incision was made either at lumbar (Th8-L1) or cervical (C2-C4) vertebra level. Using a dissecting microscope, a dorsal laminectomy of Th12 or C4 vertebra was performed to expose and cut open with a 30G needle the dura matter membrane overlying the L1-L2 or the C3-C4 spinal segments. The exposed pia matter was punctured with a 36 G penetrating needle, immediately after, a delivery needle (blunt 36 G needle) loaded with the scAAV9 vector was inserted. The precise placement of both pia-penetrating and subpial delivery needle was accomplished using a fine XYZ manipulator (SMM 100B; Narishige, Tokyo, Japan). The viral vector scAAV9-UbC-Clover-miR30-ShRNA-Control (1.0X10^13^gc/ml) or scAAV9-UbC-Clover-miR30-ShRNA-STMN2 (1.0X10^13^gc/ml) was then delivered into lumbar or cervical subpial space. The vector solution was diluted 1:2 with PBS1X just before the injection and a total of 10µl was delivered (5µl bilateral over a 5 min each) using a 50 μl Hamilton syringe and a manual infusion pump (Stoelting, cat# 51222). Finally, the delivery needle was removed, muscle was affronted, and skin was suture using 3.0 Proline. Animals received subcutaneous fluids (Ringer lactate solution) antibiotics (cefazolin, 10mg/kg) and pain medication (Buprenorphine sustain release, 0.05mg/Kg).

### Hind limb clasping Test

Clasping test has been described as a marker of disease progression in some mouse models of neurodegeneration^84, 85^. The mouse is lifted by grasping the tail near its base and maintained in this position for 10 secs. Depending on the hindlimb placement and the duration of the position the animal received a score from 0 to 3. Score of 0: the hind limbs are consistently splayed out away from the abdomen during the time the mouse is suspended; Score of 1: one hindlimb is retracted toward the abdomen for more than 50% of the time the mouse is suspended; Sore of 2: both hindlimbs are partially retracted towards the abdomen for more than 50% of the time the mouse is suspended; Score of 3: both hindlimbs are entirely retracted and touching the abdomen for more than 50% of the time the mouse is suspended. The average score of three consecutive liftings is used for each mouse.

### Grip strength test

The grip strength of forelimbs or hindlimbs paws were independently tested using a grip strength meter (Columbus Instruments, 04219). Animals were hold allowing them to grasp the bar with only the forelimbs or the hindlimbs paws and gently pulled back with steady force until both paws released the bar. Peak tension in gram was recorded for 5 consecutive trials. For germline deleted stathmin-2 mice cohorts and their matching controls, grip strength was assessed using a grid attachment to the instrument, recording the front paws and all four paws strength, the average strength in grams of three consecutive trials was recorded.

### Rotarod test

The balance and motor coordination of the mice were evaluated using the accelerating rotarod test. All the animals are subjected to two consecutive sessions, a “training session” on day one and a “test” session on day 2. On each session the mice were acclimated to the testing room for 60min. An Ugo-Basile accelerating rotarod model 47600 for mice was used. Mice were placed on a rotating rod at 4rpm which accelerates up to 40rpm over the course of 300 secs. Each mouse is subjected to four consecutive trials with a 45-sec resting interval. The time in seconds when the mouse falls from the rod is recorded. The average latency to fall of the last three trials is reported.

### Mechanical sensitivity assays

Mechanical threshold was assessed on the right and left glabrous hindpaw skin of the animals at the base of the third toe. Mice were placed into individual testing cages on a wire mesh bottom and allowed to acclimate for at least 30 minutes. Calibrated von Frey filaments (1.65–4.31) were applied to determine the 50% paw withdrawal using the Up-Down method^86–88^. Additionally, the frequency of withdrawal to supra-threshold mechanical stimuli was evaluated as a measure of mechanical sensitivity. Mice were placed into individual testing cages on a wire mesh bottom and allowed to acclimate for at least 30 minutes. Measurements were recorded by applying bending forces of 1, 4 and 8 g filaments to the base of the third toe on the plantar surface of both paws 10 times during each testing period to determine the response frequency for each filament. We chose the above Von Frey filament to warrant a supra-threshold stimulus within the noxious mechanical range^89–92^. The average ± standard error of the withdrawal response frequency for each filament force /paw was recorded and plotted. This method was adapted from previously studies^93–95^.

### Compound muscle action potential

Compound action potential (CMAP) analysis was conducted as previously described^96, 97^. Briefly, mice were anesthetized with isoflurane (1.5–2%) and placed on a thermostatically regulated heating pad to maintain normal body temperature, CMAP responses were recorded from the proximal hindlimb using two recording electrodes. The active electrode was positioned over the tibialis anterior (TA) muscle. The reference electrode was positioned at the metatarsal region of the foot on the same limb. Supramaximal stimulation of the sciatic nerve was elicited via two needle electrodes placed subcutaneously. Over the sacrum (anode) and the sciatic notch (cathode). Stimulations were delivered through a Stimulus Isolator (FE180, AD Instruments). CMAP amplitude was recorded and the peak-to-peak amplitude of the CMAPs was measured. For mice with stathmin-2 knock-down either by ASOs or subpially delivered AAV9, responses were recorded from the TA using 30G platinum transcutaneous needle electrodes (distance between recording electrodes ∼1cm; Grass Technologies, Astro-Med). Recording electrodes were connected to an active headstage (3110W; Warner Instruments) amplified signal was acquired by the PowerLab 8/30 data acquisition system (ADInstruments) at a sampling frequency of 20kHz, digitized and analyzed.

### Nerve conduction velocity

Nerve conduction velocity at the sciatic nerve was previously described^98^. It was determined by measuring differences in compound muscle action potential (CMAP) latency in the muscles of the hind paw following sciatic nerve stimulation at two points, hip and ankle. Briefly, mice were anesthetized with isoflurane (1.5 – 2 %) and placed on a thermostatically regulated heating pad to maintain normal body temperature. For recording, the active needle electrode was inserted in the center of the paw and a reference electrode was placed in the skin between the first and second digits. The distance between points of stimulation and the recorded latencies was used in the calculation of velocity.

### Tissue collection

Animals were euthanized with pentobarbital (100mg/kg) and perfused with 20 ml of ice-cold phosphate buffered saline (PBS 1X). The mouse tissues were dissected and collected as follow: the spinal cord was divided in cervical, thoracic, and lumbar segments. Each segment was then divided in 3 equal size pieces. The rostral pieces were post-fix for 24 h in freshly prepared 4% PFA (4% paraformaldehyde dissolved in 0.15 M sodium phosphate buffer, pH 7.4) at 4 °C and the other 2 pieces were flash frozen in dry ice and saved for qPCR and Western blot analysis. Left and right muscles (gastrocnemius, tibialis, peroneal and soleus) were dissected and post-fix for 5 days in 4% PFA at 4 °C. The right and left dorsal root ganglion L5 (DRG L5) were dissected and post-fixed for 24 h in 4% PFA at 4 °C, the remaining right and left lumbar DRG’s (L3,4, 6) were collected as a pool and flash frozen in dry ice for qPCR analysis. All designated samples for immunohistochemistry or FISH were cryo-protected for 72 hours in 30% phosphate buffer after their post-fixation period and stored at 4 °C until processing. The right sciatic nerve and the dorsal and ventral roots of left and right DRG L5 were dissected and kept in freshly prepared 4% PFA until further processing for axonal counting. The left sciatic nerve and the liver were flash frozen for western blot analysis.

Animals used for electron microscopy were euthanized with pentobarbital and the whole body perfused with 20 ml of ice-cold PBS 1x followed by 20 ml freshly prepared fixative (2 % PFA, 2 % glutaraldehyde, 0.1 M cacodylate buffer pH=7.2). After perfusion DRG’s with their dorsal and ventral roots from L5 and sciatic nerves were dissected, dropped in the same fixative solution, and kept at 4 °C until processing.

### Immunohistochemistry

Cryoprotected tissues (spinal cords, muscles and DRGs) were embedded in Optimal Cutting Temperature (OCT) matrix compound (Tissue-Tek, Sakura Finetek), frozen with dry ice and mounted in the cryostat. Sections were cut from the spinal cords and muscles (30-μm thickness), or dorsal root ganglion (15-μm thickness). Free-floating sections were washed three times in PBS 1X with 0.3% Triton-X100 followed by a blocking step with 5% donkey serum in PBS 1X with 0.3% Triton-X100 for 1h. Sections were then incubated in the primary antibodies (in blocking solution) overnight at 4 °C. Next day, the sections were washed three times with PBS 1X with 0.3% Triton-X100, and incubated with secondary antibody in PBS, 0.3% Triton-X100 for 1h at room temperature. Sections were mounted on slides, dried at room temperature and cover slipped in ProLong Gold antifade mounting medium, with DAPI (Invitrogen). The primary and secondary antibodies and their dilutions are described on Table 2. Antigen retrieval was needed to obtain a signal for Stmn2 protein in the spinal cord and DRG. Briefly, thirty-micron sections were mounted on slides and let dried overnight. A hydrophobic ring was then drawn around the sections and they were washed 3 times with PBS 1X with 0.3% Triton-X100. Immediately after the slides were immersed in pre-heated (90-100 degree Celsius) 1X antigen retrieval solution (Dako, cat # S1699) for 25 min, removed and washed with PBS 1X.

**Table 2.** Antibodies.

| Primary Antibodies |  |  |  |  |  |
| --- | --- | --- | --- | --- | --- |
| Name | Cat # | Company | Host | Dilution | Application |
| NeuN | ABN78 | Millipore-Sigma | rabbit | 1:1000 | IHC |
| CHAT (Choline acetyltransferase) | AB144P | Millipore-Sigma | goat | 1:100 | IHC |
| GFAP-Cy3 conjugated (clone G-A-5) | C9205 | Millipore-Sigma | mouse | 1:1000 | IHC |
| Iba1 (clone NCNP24) | 019-19,741 | Wako | rabbit | 1:1000 | IHC |
| α-BTX (α-bungarotoxin, Alexa Fluor 555 conjugated) | B35451 | Invitrogen | ----- | 1:500 | IHC |
| Synaptophysin | NBP2-25170 | Novus Biologicals | chicken | 1:500 | IHC |
| Neurofilament-H | AB5539 | Millipore-Sigma | chicken | 1:500 | IHC |
| Calcitonin gene related peptide (CGRP) | Ab36001 | Abcam | goat | 1:1000 | IHC |
| Lectin IB4 conjugated to Alexa- Fluor 647 (anti-IB4 Alexa Fluor 647) | I32450 | Molecular Probes | ----- | 1:2000 | IHC |
| Human/Mouse Stathmin-2/STMN2 Clone #684433 | MAB6930 | R&D | mouse | 1:200 | IHC (antigen retrieval) |
| Anti-Neurofilament-H (SMI-31) | 50-102-9514 | Biolegend | mouse | 1:1000 | WB |
| Neurofilament-H Clone RT-97 | NBP2-29435 | Novus Biologicals | mouse | 1:500 | WB |
| NF-M, Clone NN18 | N5264 | Sigma | mouse | 1:1000 | WB |
| NF-L, Clone NR4 | N5139 | Sigma | mouse | 1:1000 | WB |
| Mouse Tubulin β3, Clone TUJ1; | 801202 | Biolegend | mouse | 1:4000 | WB |
| Rabbit Tubulin β3 Clone TUJ1; | 845502 | Biolegend | rabbit | 1:4000 | WB |
| Stathmin-2 | 10586-1-AP | Proteintech | rabbit | 1:1000 | WB |
| Stathmin-2 | NBP1-49461 | Novus Biologicals | rabbit | 1:1000 | WB |
| GAPDH Clone 6C5 | ab8245 | Abcam | mouse | 1:5000 | WB |
| GAPDH Clone 14C10 | 2118 | Cell Signaling | rabbit | 1:5000 | WB |
| **Secondary Antibodies** |  |  |  |  |  |
| **Name** | **Cat #** | **Company** | **Host** | **Dilution** | **Application** |
| Cy3 AffiniPure Donkey anti-Guinea Pig IgG [H1L] | 706-165-148 | Jackson ImmunoResearch | Donkey | 1:1000 | IHC |
| Alexa 488 Donkey anti-Goat IgG [H1L] | A-11,055 | Invitrogen | Donkey | 1:1000 | IHC |
| Alexa 549 Donkey anti-Rabbit IgG [H1L] | A-21,207 | Invitrogen | Donkey | 1:1000 | IHC |

### Fluorescent In situ Hybridization - FISH

A paired double-Z oligonucleotide probe against Mm-stmn2 RNA pre-designed and commercially available from RNAscope (498391-C2; NM_025285.2, 20 pairs, 898 -1849 nucleotides) was obtained (Table 3). Pre-fixed tissue was embedded in OCT (Tissue-Tek Sakura Finetek), frozen in dry ice and mounted in the cryostat. Sections from the spinal cord and DRGs were cut (thickness of 15 µm), directly mounted on Superfrost Plus slides (Thermo Fisher) and left overnight at room temperature to dry. The following day, RNA FISH was performed using the RNAscope Multiplex Fluorescent v.2 (no. 323100), following the fixed frozen tissue protocol according to the manufacturer’s instructions. Briefly, the tissues were treated with peroxidase hydrogen blocker before boiling at 98– 100 °C in a pretreatment solution for 10 min. Protease plus was then applied for 30 min at 40 °C. The target probe (RNAscope® Probe-Mm-Stmn2-C2) was prediluted 1:50 and hybridized for 2 h at 40 °C, followed by a series of signal amplification and washing steps. All incubation steps at 40 °C were performed in a HybEZ Hybridization System. Hybridization signals were detected by a chromogenic reaction using red chromogen dilution 1:3000 (PerkinElmer TSA Plus Cyanine 3 System). RNA-staining signal was identified as red punctate dots and clusters. RNA quality was evaluated for each sample using RNAscope 3-plex Positive Control Probe mouse specific containing the housekeeping gene cyclophilin B (PPIB), RNA polymerase subunit IIA (PolR2A) and ubiquitin C (UBC). Negative control background staining was evaluated using a probe specific to the bacterial dapB gene (RNAscope 3-plex Negative Control Probe).

**Table 3.**
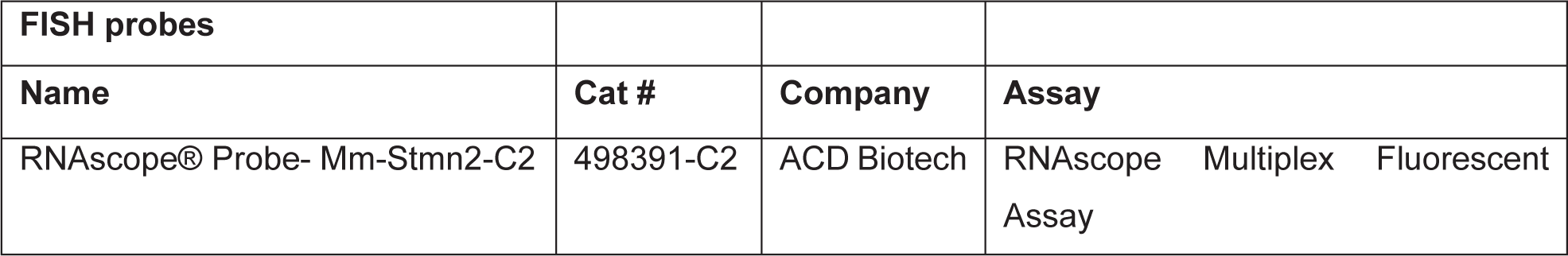
FISH probes.

| **FISH probes** |  |  |  |
| --- | --- | --- | --- |
| **Name** | **Cat #** | **Company** | **Assay** |
| RNAscope® Probe- Mm-Stmn2-C2 | 498391-C2 | ACD Biotech | RNAscope Multiplex Fluorescent Assay |

### Immunofluorescence

Mice were perfused intracardially and fixed with 4% paraformaldehyde in 0.1 M Sorenson’s phosphate buffer, pH 7.2, the tissue was dissected, post-fixed for 2 hours in the same fixative and transferred in a 30% sucrose phosphate buffer for at least 2 days. The lumbar spinal cords and brains were embedded in OCT compound (Sakura) and snap frozen in isopentane (2-methylbutane) cooled at 40 C on dry ice. Floating lumbar spinal cord or brain cryosections (30mm or 35mm, respectively) were incubated in a blocking solution containing PBS1x, 0.5% Tween-20, 1.5% BSA for 1.5 hours at room temperature and then in PBS1x, 0.3% Triton X-100 overnight at room temperature with the primary antibodies (Table 2). Primary antibodies were washed with PBS1x and then detected using donkey anti-rabbit or anti-mouse FITC or Cy3 (1:500) coupled secondary antibodies (Jackson ImmunoResearch). The sections were washed with PBS1x and mounted. Analysis was performed on Zeiss SP8 confocal microscope. Fluorescence intensity from unsaturated images captured with identical confocal settings.

### Motor neuron counting

Lumbar ChAT positive ventral horn motor neurons were counted from 10-15 lumbar spinal cord cryosections (per animal) spaced 300μm apart and expressed as total motor neurons counted per spinal cord hemi section.

### Neuromuscular junction innervation

Gastrocnemius muscle was dissected from perfused mice and prepared as described in the immunofluorescence section. Floating 40 μm thick longitudinal sections of gastrocnemius were incubated in a blocking solution containing PBS1x, 0.5% Tween-20, 1.5% BSA for 4 hours at room temperature and then in PBS1x, 0.3% Triton X-100 overnight at room temperature with the polyclonal rabbit anti-synaptophysin antibody at 1:50 (Invitrogen). The sections were washed with PBS1x and then incubated with donkey antirabbit Cy3 (Jackson ImmunoResearch) and a-bungarotoxin-Alexa488 (Invitrogen) at 1:500 for 2 hours at room temperature. The sections were further washed with PBS1x and mounted. Analysis was performed on SP8 Leica confocal microscope. Individual NMJs were considered as innervated when the presynaptic axon, (labelled either by neurofilament-H+synaptophysin or β3-tubulin+synaptophysin) staining covered at least 50% of the area of α-bungarotoxin staining.

### CGRP and IB4 Quantification in the spinal cord

The experimenter was blind during the image analysis, and the order of images analyzed was randomized. A maximum projection of all the z-stacks was obtained for analysis using Fiji Software^99^. A region of interest was created to selectively analyze CGRP and IB4 projections arriving to the dorsal horn of the spinal cord. To enhance detection of the borders, images were blurred with a gaussian blur. After, the area of CGRP and IB4 staining was determined by counting the pixels with intensity higher than a pre-set threshold (constant for all analyzed images) inside the respective regions of interest.

### Morphometric Analysis of Axons

Mice were perfused intracardially and fixed with 4% paraformaldehyde in 0.1 M Sorenson’s phosphate buffer, pH 7.2, and the L5 lumbar roots were dissected and conserved at 4 C in a 2 % PFA, 2 % glutaraldehyde, 0.1 M cacodylate buffer (pH=7.2). L5 roots were embedded in Epon-Araldite as described in the electron microscopy section, thick sections (0.75um) were prepared and stained for light microscopy with toluidine blue. Cross sections of L5 motor axons were analyzed at each age group. Axonal diameters were measured using ImageJ and the diameters of all axons in the ventral and dorsal roots were determined.

### RNA extraction for qRT-PCR quantification

Tissue sections snap frozen at the time of collection were mechanically homogenized in 1ml Trizol (Thermo) with a rotor stator homogenizer (Omni International) and RNA was chloroform extracted as per the manufacturer protocol. RNA was quantified on a Nanodrop spectrophotometer (Thermo) and 1μg taken forward to first-strand cDNA synthesis with SuperScript-III reverse transcriptase (Thermo) and oligo dT priming. For DRGs, where total recoverable RNA is more limiting, 200ng of RNA was utilized for first-strand cDNA synthesis. cDNA was diluted to 1ng/μL, and 4ng loaded per 10μL SYBR Green reaction (BioRad), with three technical replicates loaded per biological sample, run on a C1000 thermocycler with a 384-well qPCR reaction module (BioRad) using primers detailed in Table 2. All melt-curves showed a single distinct amplification product. Murine GAPDH and RPS9 genes were used as endogenous controls and showed equivalent results. The data normalized with GAPDH was graphed for the figures. Relative expression for each gene was calculated from delta-Cq data, with graphing and statistical analysis performed in Prism8 software (GraphPad). Data collection and analysis was performed blinded to the conditions of the experiments.

### Tissue Protein Extraction and Immunoblotting

Mouse spinal cords and sciatic nerves were homogenized in cold RIPA Buffer supplemented with protease and phosphatase inhibitors and extracts were clarified by cold centrifugation for 20 minutes at 20,000g. Equal protein amounts were separated by SDS-PAGE, transferred to nitrocellulose membranes, and probed with the indicated antibodies diluted in 5% nonfat dry milk in TBS-Tween 0.1%, followed by horseradish peroxidase-conjugates secondary antibodies (Jackson ImmunoResearch). Immunoblots were developed on film through ECL-mediated chemiluminescence.

### Statistical analysis

Data were analyzed and graphed using Prism 9.02 software (GraphPad, San Diego, CA). Mann-Whitney t-test was used for comparisons between two groups. One-way or Two-way ANOVA test with Dunnett’s correction was used for multiple group comparison. Statistical significance was set at p<0.05 for all comparisons.

## Acknowledgements

We are grateful to the Muscular Dystrophy Association for supporting JL-E and ZM with MDA Development grants. We thank UCSD Genetics Training Program and the National Institute for General Medical Sciences, T32 GM008666 for supporting MWB and T32AG066596-01 for supporting MSB. This work was supported by grants from ALS finding a Cure (to CL-T), NINDS/NIH: R01NS112503 to MM, CL-T and DWC, RF1NS124203 to CML, DWC and CL-T. The microscope core was supported by NINDS/NIH P30NS047101. CL-T is the recipient of the Araminta Broch-Healey Endowed Chair in ALS.

---

**Supplementary Figure 1:**
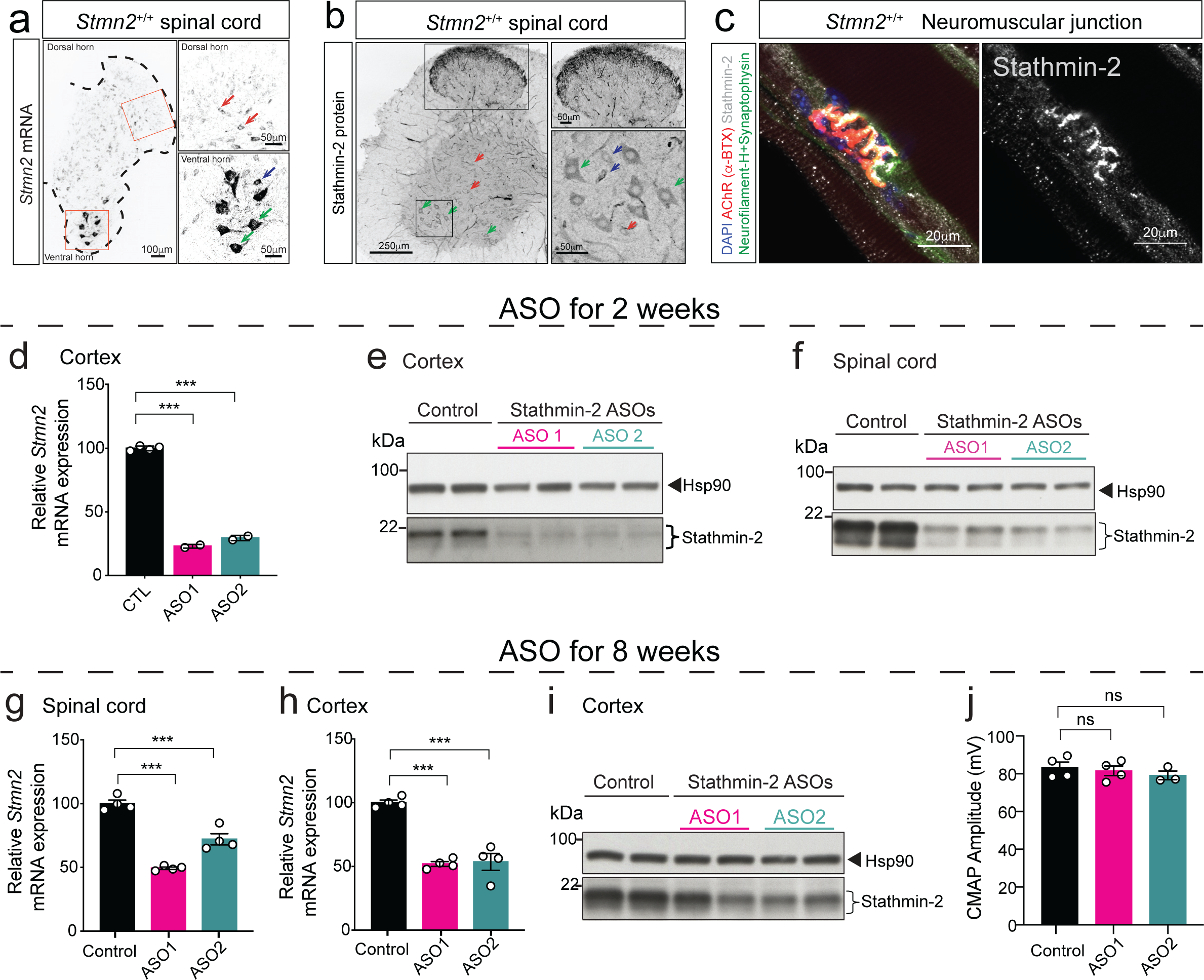
Intraventricular ASO delivery efficiently reduces stathmin-2 expression in mouse cortex and spinal cord. **(a)** *Stmn2* mRNA levels detected by FISH in lumbar spinal cord hemisections from 12-month-old WT mice. Green arrows: *α*-motor neurons; blue arrows: *γ*-motor neurons; red arrows: interneurons. **(b)** Immunofluorescence confocal image of 12-month-old WT mice spinal cord hemisection immunolabeled for stathmin-2 protein. Green arrows: *α*-motor neurons; blue arrows: *γ*-motor neurons; red arrows: interneurons. **(c)** Immunofluorescence confocal image of 12-month-old WT mice gastrocnemius muscle revealing stathmin-2 presence at the neuromuscular junction. **(d,e)** Quantification of *Stmn2* mRNA levels by qPCR **(d)** and immunoblots **(e)** showing stathmin-2 protein levels in mice cortex 2 weeks after the ICV injection of non-targeting or *Stmn2* targeting ASOs. Hsp90 was used as a loading control in the immunoblotting. **(f)** Immunoblots showing stathmin-2 protein levels in mouse spinal cord 2 weeks after the ICV injection of non-targeting or *Stmn2* targeting ASOs. Hsp90 was used as a loading control. *Indicates non-specific band. **(g)** Quantification of *Stmn2* mRNA levels by qPCR in mouse spinal cord 8 weeks after the ICV injection of non-targeting or *Stmn2* targeting ASOs. *Gapdh* was used as an endogenous control gene. Each datapoint represents an individual mouse. Error bars are plotted as SEM. **(h,i)** Quantification of *Stmn2* mRNA levels by qPCR **(h)** and immunoblots **(i)** showing stathmin-2 protein levels in mice cortex 8 weeks after the ICV injection of non-targeting or *Stmn2* targeting ASOs. Hsp90 was used as a loading control in the immunoblotting. *Gapdh* was used as an endogenous control gene. Each datapoint represents an individual mouse. Error bars are plotted as SEM. **(j)** Compound muscle action potential (CMAP) measurements in muscles of WT mice treated with non-targeting or *Stmn2* targeting ASOs for 8 weeks.

**Supplementary Figure 2:**
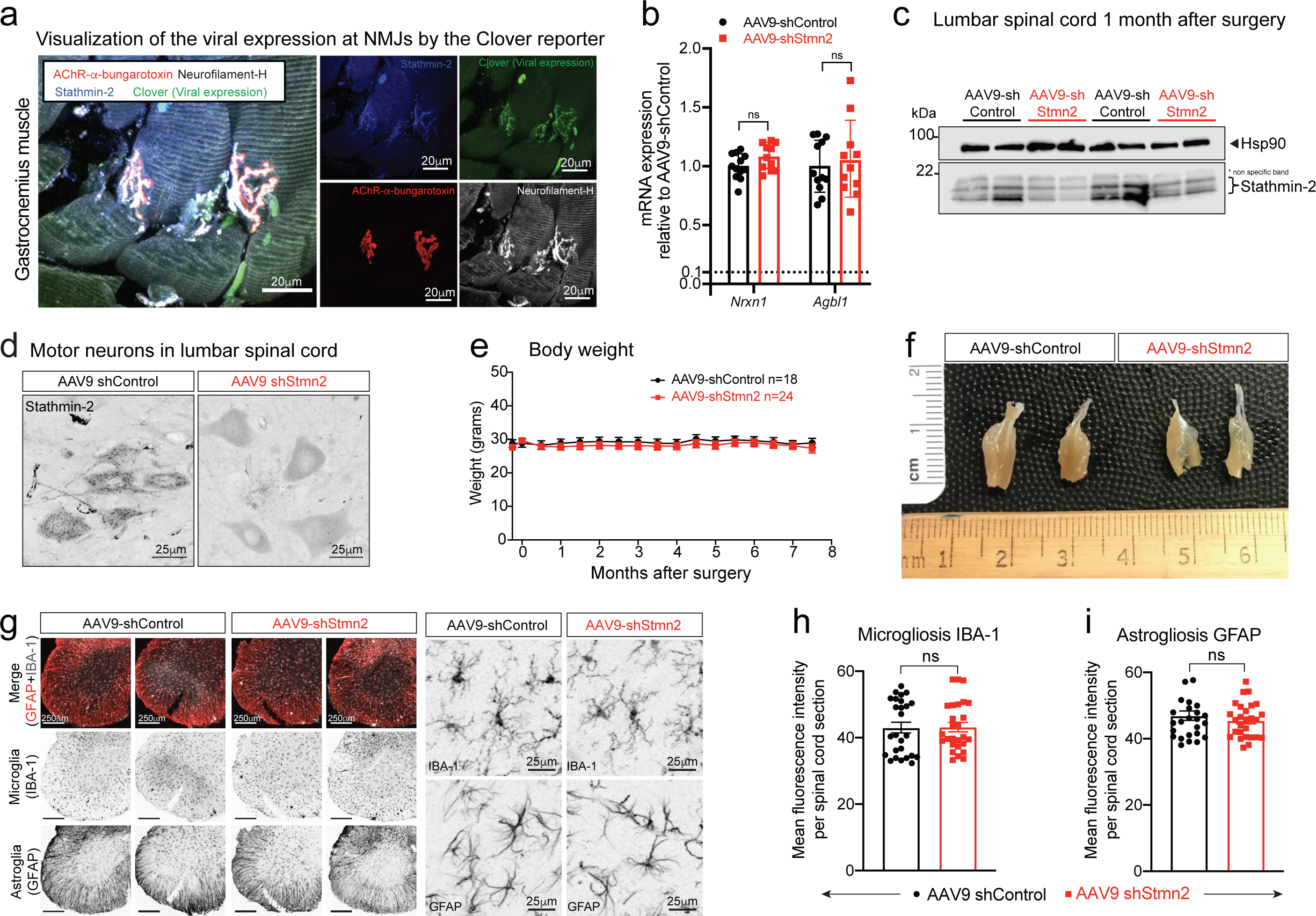
Sustained stathmin-2 depletion induces axonal withdrawal from neuromuscular junctions without compromising motor neuron survival. **(a)** Representative confocal image of gastrocnemius muscle stained for stathmin-2 (blue), AChR cluster in the muscle using *α*-bungarotoxin (red), direct imaging of clover in the 488-wavelength representing viral expression and neurofilament-H in white. **(b)** Measurement in lumbar spinal cord segments at 8-months post injection of control or *Stmn2* targeting AAV9 of potential off-target genes mRNA expression by qRT-PCR confirming selective *Stmn2* suppression using the designed RNAi strategy. *Gapdh* was used as an endogenous control gene. Each data point represents an individual mouse. Error bars are plotted as SEM. **(c)** Immunoblots to determine stathmin-2 protein level in mouse lumbar spinal cord 1-month after subpial injection of a *Stmn2* reducing AAV9 or after injection of a non-targeting control AAV9. Hsp90 was used as a loading control. *Indicates non-specific band. **(d)** Mouse lumbar spinal cord immunofluorescence micrographs visualized with stathmin-2 antibody 8 months after subpial injection into the lumbar spinal cord of non-targeting control AAV9 or *Stmn2-*reducing AAV9. **(e)** Bi-weekly measurements of mouse body weight after subpial injection of AAV9 encoding either non-targeting sequence or shRNA against *Stmn2*. **(f)** Representative images of entire gastrocnemius muscles from mice 8 months after subpial delivery of AAV9 encoding either a non-targeting control AAV9 or an AAV9 encoding an shRNA against *Stmn2*. **(g-i)** Representative immunofluorescence images of mouse lumbar spinal cord stained with the microglial and astrocytic markers IBA1 and GFAP **(g)** and quantification of microgliosis **(h)** and astrogliosis **(i)**, 8 months after subpial delivery of a non-targeting control AAV9 or an AAV9 encoding an shRNA against *Stmn2*.

**Supplementary Figure 3:**
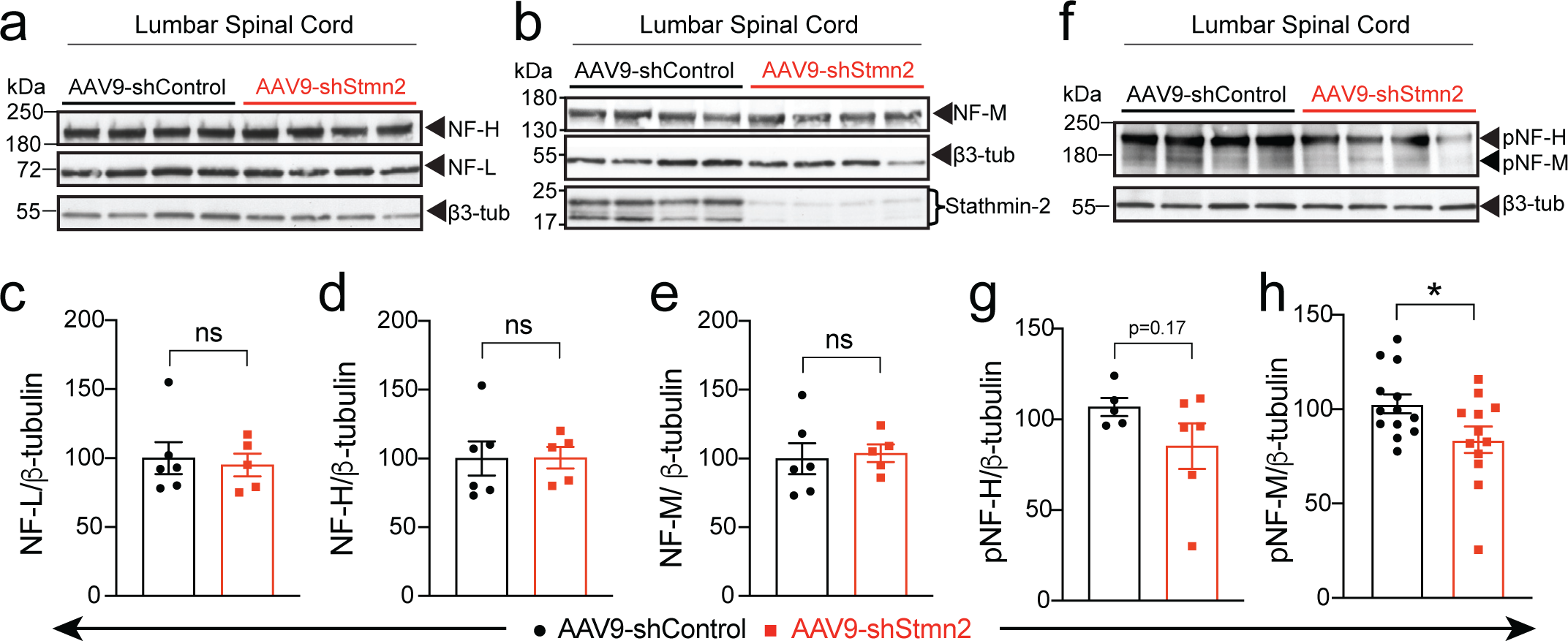
Decreased phosphorylation of NF-M and NF-H upon sustained *Stmn2* suppression. **(a-b)** Levels of non-phosphorylated neurofilament H (NF-H) and levels of neurofilament light (NF-L) **(a)** and total neurofilament medium (NF-M) **(b)** analyzed by immunoblotting spinal cord extracts from WT mice 8 months after subpial injection of either AAV9 encoding a non-targeting sequence or encoding an shRNA against *Stmn2*. β3-tubulin was used as loading control. AAV9-shRNA-mediated suppression of stathmin-2 protein levels was confirmed in all examined samples **(b)**. **(c-e)** Quantification of the immunoblots in panels a,b. Each point represents an individual animal. Error bars are plotted as SEM. **(f)** Levels of phosphorylated neurofilament heavy (pNF-H) and medium (pNF-M) subunits analyzed by immunoblotting of spinal cord extracts from WT mice, 8 months after subpially injected either with AAV9 encoding a non-targeting sequence or encoding an shRNA against *Stmn2*. β3-tubulin was used as a loading control. **(g,h)** Quantification of the immunoblots from panel f. Each point represents an individual animal. Error bars are plotted as SEM.

**Supplementary Figure 4:**
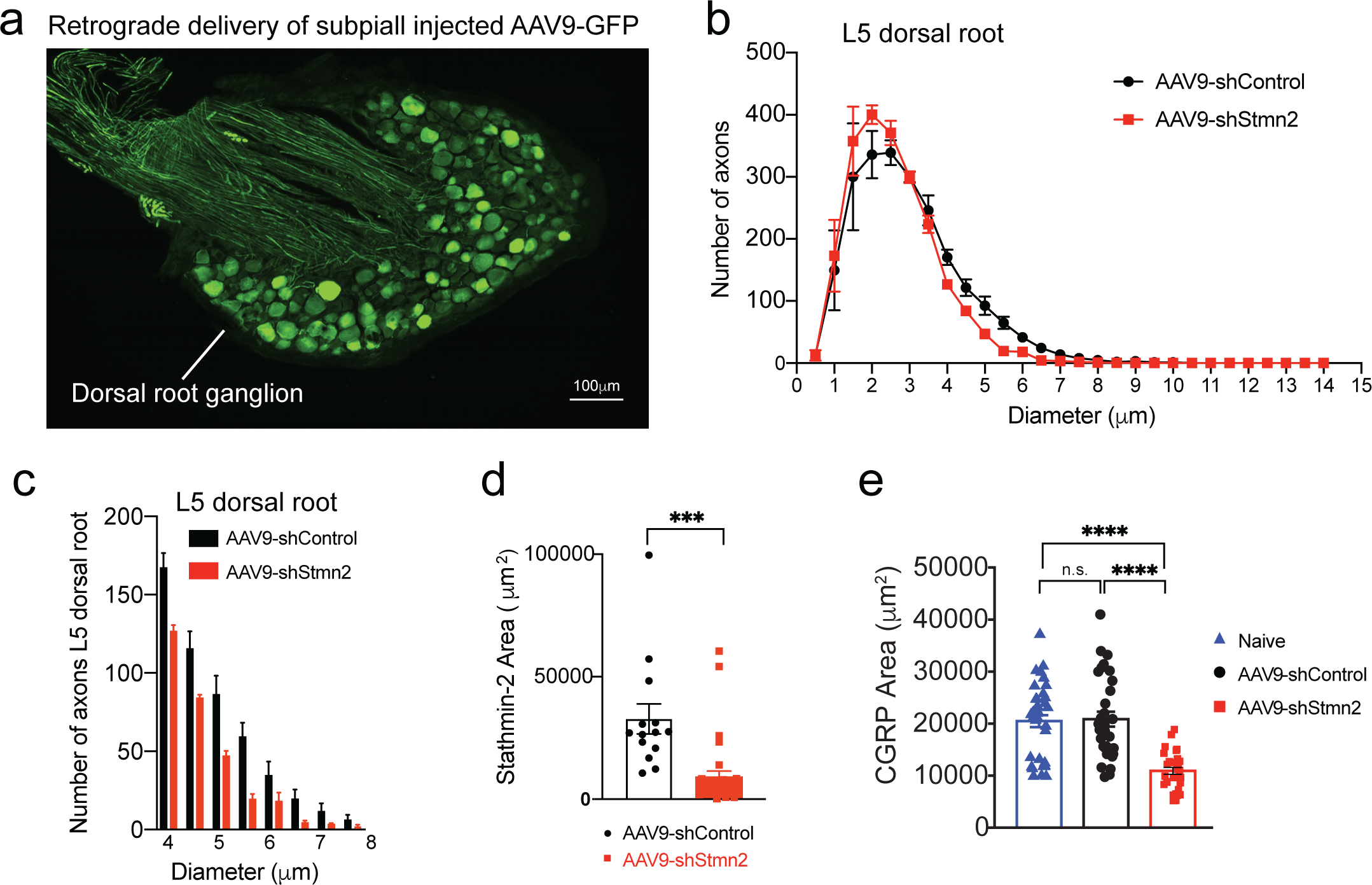
Reduced stathmin-2 levels by subpial injection alters sensory marker in lumbar spinal cord. **(a)** Representative image of lumbar dorsal root ganglion 2 months after subpial injection into lumbar spinal cord and subsequent retrograde delivery of AAV9 expressing green fluorescent protein (GFP). **(b)** Size distribution of axonal diameter of sensory neurons innervating the dorsal spinal cord. Error bars are plotted as SEM. **(c)** Axon numbers in the 4 μm to 8 μm diameter range in the L5 dorsal root. **(d-e)** Quantification related to Figure 7i,k of positive area for stathmin-2 **(d)** and CGRP **(e)** in the dorsal spinal cord 8 months after subpial delivery of AAV9 encoding either non-targeting sequence or sh*Stmn2*.

**Supplementary Figure 5:**
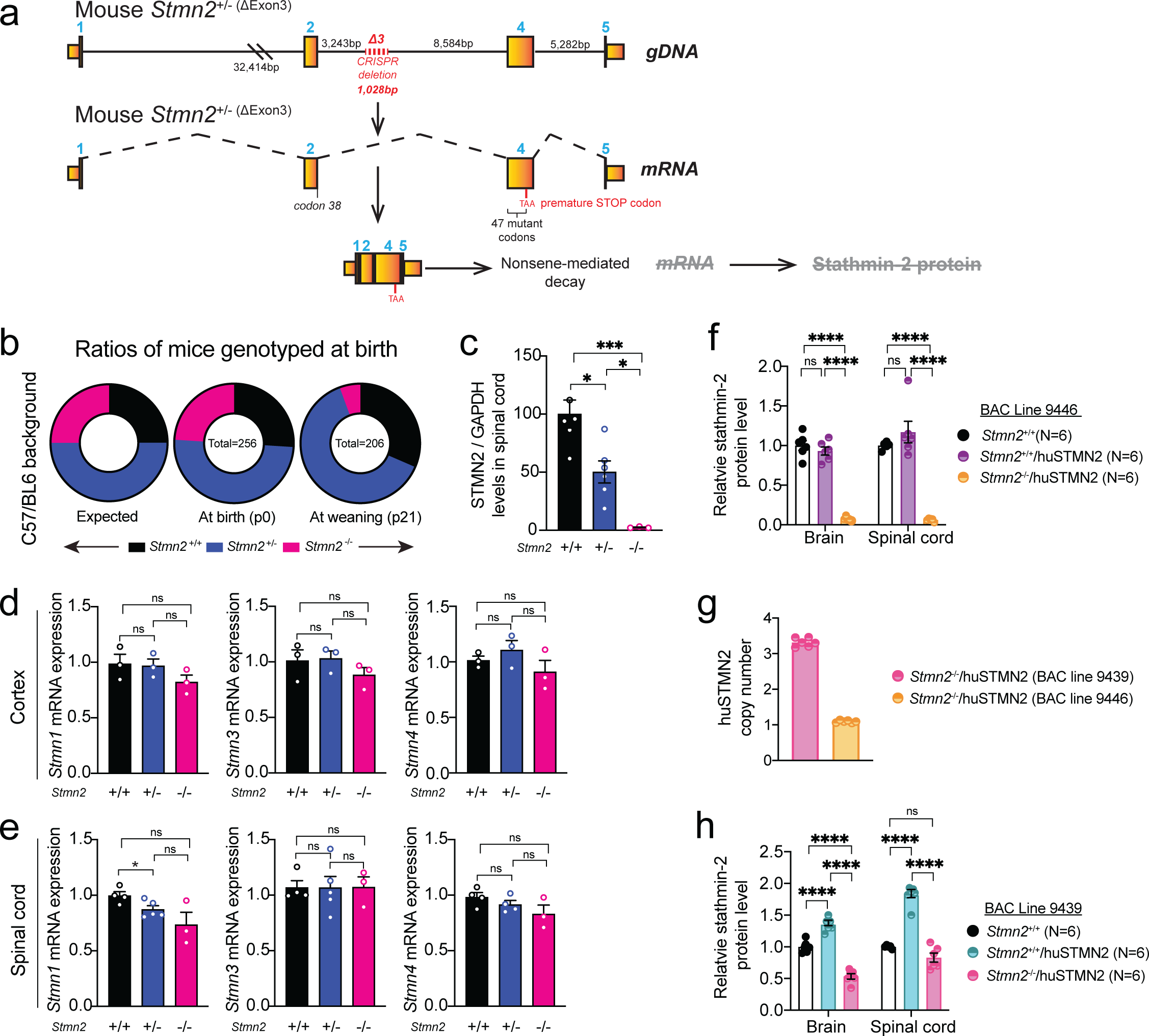
Stathmin-2 related genes remain unchanged upon stathmin-2 loss. **(a)** Diagram of genome editing design by CRISPR-Cas9-mediated excision of mouse *Stmn2* exon 3 that leads to complete absence of stathmin-2 protein. **(b)** Ratios of mice expected, genotyped at birth (p0) and alive at weaning age (p21) from *Stmn2*^+/-^ to *Stmn2*^+/-^ crossing in C57/Bl6 background. **(c)** Stathmin-2 protein quantification from the immunoblots in Fig. 1e normalized by GAPDH. **(d,e)** Measurement of mouse *Stmn*-1, -3 and -4 mRNA levels extracted from 12-month-old cortex **(d)** and spinal cord **(e)** of *Stmn2*^+/+^, *Stmn2*^+/-^ *Stmn2*^-/-^ mice. *Gapdh* was used as an endogenous control gene. Each data point represents an individual mouse. Error bars are plotted as SEM. Statistical analysis by Kruskal Wallis nonparametric tests *p<0.05 **p<0.01 ***p<0.001. **(f)** Stathmin-2 protein quantification from brain and spinal cord extract of *Stmn2*^+/+^, *Stmn2*^+/+^/huSTMN2 and *Stmn2*^-/-^/huSTMN2 (BAC line 9439) by immunoblotting. **(g)** huSTMN2 transgene copy numbers measured in BAC transgenic lines 9439 and 9446. **(h)** Stathmin-2 protein quantification from brain and spinal cord extract of *Stmn2*^+/+^, *Stmn2*^+/+^/huSTMN2 and in *Stmn2*^-/-^/huSTMN2 (BAC line 9446) by immunoblotting.

**Supplementary Figure 6:**
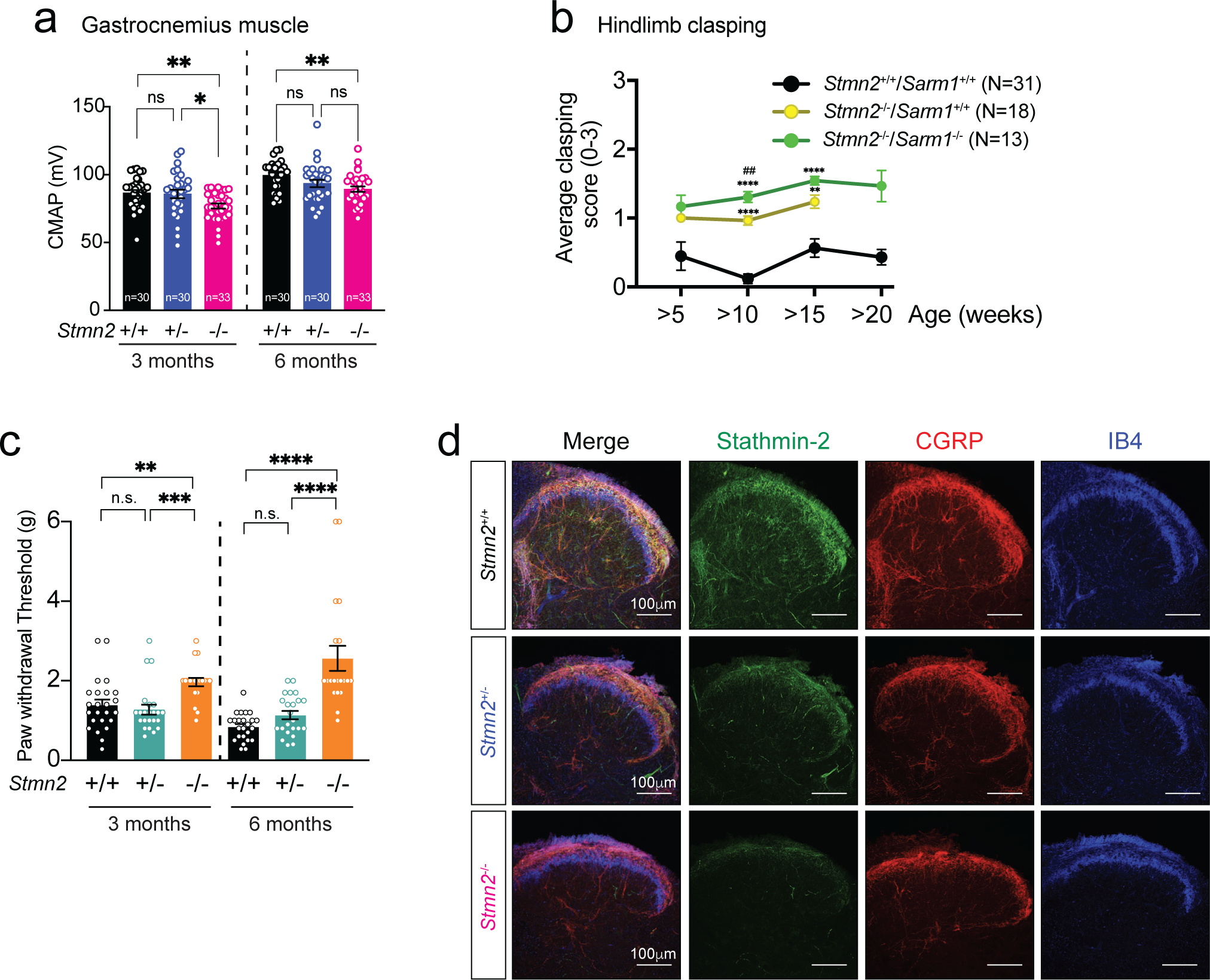
Absence of stathmin-2 results in motor deficits regardless of the genetic background. **(a)** Compound muscle action potential (CMAP) measurements in gastrocnemius muscle of *Stmn2*^+/+^ and *Stmn2*^+/-^ and *Stmn2^-/-^* mice at 3 and 6 months of age. **(b)** Hindlimb clasping test of *Stmn2*^+/+^/*Sarm1*^+/+^, *Stmn2*^+/+^/*Sarm1*^-/-^ and *Stmn2*^-/-^/*Sarm1*^-/-^ mice in C57/BL6 background for 20 weeks. Statistical analysis by Tukeýs multiple comparison test **p<0.01; ****p<0.0001 when comparing to *Stmn2*^+/+^/*Sarm1*^+/+^. ^##^p< 0.01 when comparing between *Stmn2*^+/+^/*Sarm1*^-/-^ and *Stmn2*^-/-^/*Sarm1*^-/-^. **(c)** Von Frey analysis for the sensory response in hindlimbs of *Stmn2*^+/+^, *Stmn2*^+/-^ and *Stmn2*^-/-^ mice in FVB background at 6 months of age. **(d)** Lumbar spinal cord dorsal section of *Stmn2*^+/+^, *Stmn2*^+/-^ and *Stmn2*^-/-^ mice immunostained for stathmin-2 (green), CGRP (red) and Isolectin B4 (IB4) in blue.

**Supplementary Figure 7:**
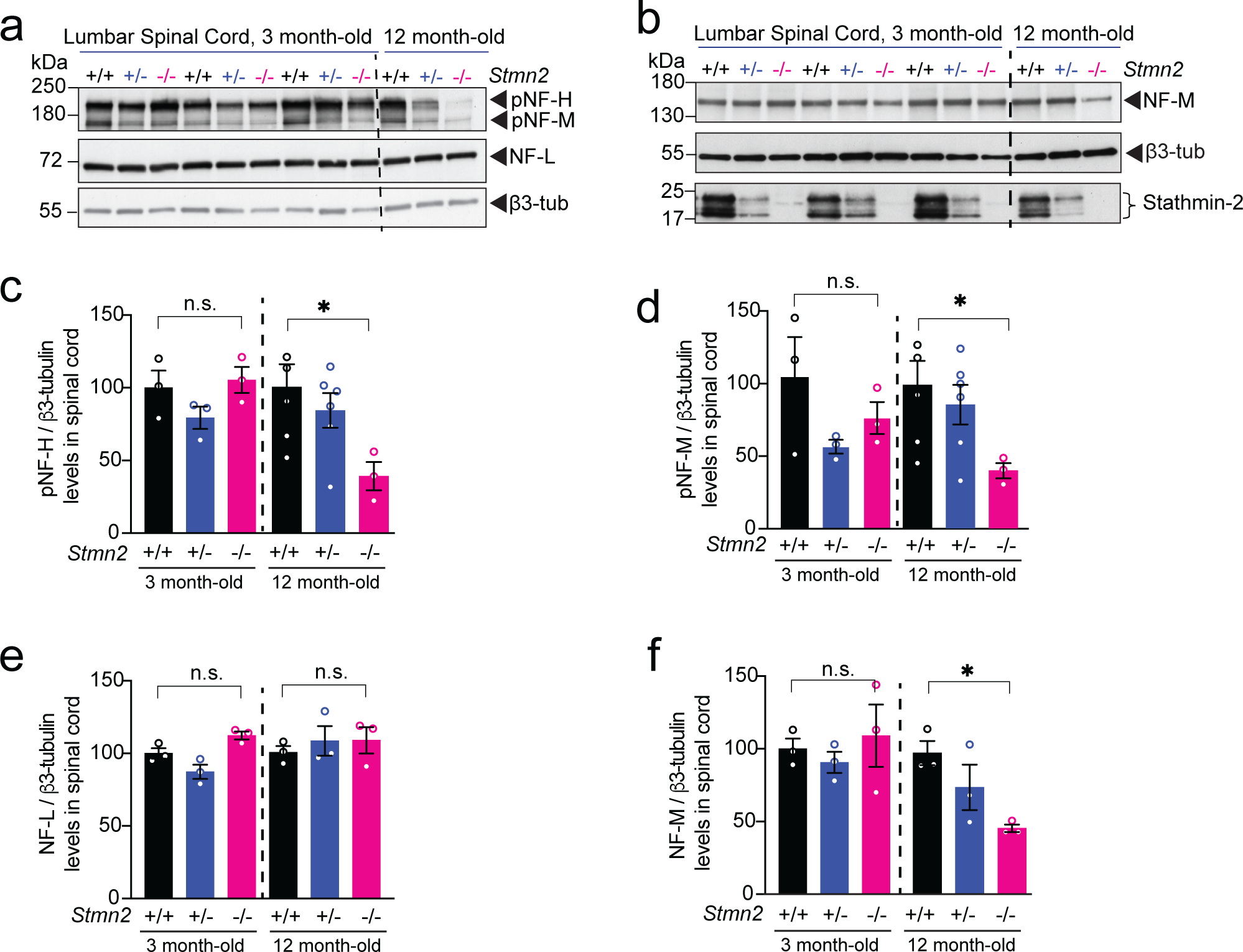
Absence of stathmin-2 alters neurofilament composition over time in mice spinal cords. **(a)** Immunoblotting for phosphorylated neurofilament heavy (pNF-H), phosphorylated neurofilament medium (pNF-M), and neurofilament light (NF-L) analyzed on 3 and 12 months-old *Stmn2*^+/+^, *Stmn2*^+/-^ and *Stmn2*^-/-^ mice lumbar spinal cord protein extracts. **(b)** Immunoblotting for neurofilaments medium (NF-M) and stathmin-2 from *Stmn2*^+/+^, *Stmn2*^+/-^ and *Stmn2*^-/-^ mice lumbar spinal cord protein extracts at 3 and 12 months-old. β3-tubulin was used as protein loading control. Stathmin-2 levels of *Stmn2*^+/+^, *Stmn2*^+/-^ and *Stmn2*^-/-^ at ∼21 kDa are also shown. **(c-f)** Quantifications from immunoblots for pNF-H **(c)**, NF-M **(d)**, NF-L **(e)** and NF-M **(f)** are shown. β3-tubulin remained unchanged confirming amount of protein loading control.

